# MYC/MIZ1 suppression of lysosomal protein degradation drives immune evasion in pancreatic ductal adenocarcinoma

**DOI:** 10.64898/2026.07.23.740267

**Authors:** Bastian Krenz, Toshitha Kannan, Greta Mattavelli, Werner Schmitz, Anneli Gebhardt-Wolf, Mara John, Ana Cetkovic, Arpa Aintablian, Stella Caviccioli, Lukas Nowak, Sarah Bopp, Christina Schülein-Völk, Carsten P. Ade, Steffi Herold, Julian Bender, Bettina Warscheid, Christine Sers, Magdalena Zukowska, Martin Vaeth, Gabriele Büchel, Frederique Zindy, Shondra M. Pruett-Miller, Martine F. Roussel, Dieter Saur, Angela Riedel, Martin Eilers

## Abstract

The MYC oncoprotein promotes immune evasion of pancreatic ductal adenocarcinoma (PDAC), but the underlying molecular mechanisms are not fully understood. Here we show that MYC protects PDAC tumors from CD4^+^ T cell-dependent elimination. Single cell sequencing shows that MYC suppression in tumor cells increases amino acid availability and broadly activates amino acid-responsive gene expression programs in immune cell populations. This occurs because MYC-driven uptake depletes free amino acids from tumor interstitial fluid and plasma, while MYC compromises macropinocytosis and autophagy, both of which depend on lysosomal protein degradation. MYC engages the POZ/BTB transcription factor MIZ1 to suppress lysosomal genes regulated by the TFE3/TFEB/MITF network or by free MIZ1, thereby inhibiting lysosomal protein degradation. An orthogonal genetic model enabling transient, selective inhibition of amino acid uptake in tumor cells recapitulates the effects of MYC depletion on amino acid levels in the tumor microenvironment and induces complete, CD4^+^ T cell-dependent tumor eradication with long-term survival. We propose that MYC-mediated, cell-autonomous disruption of lysosome function coupled to non-cell-autonomous protection from immune clearance allows MYC-low cells to benefit from MYC-high neighbors, such that intratumoral heterogeneity in MYC expression confers a selective advantage to the entire tumor.

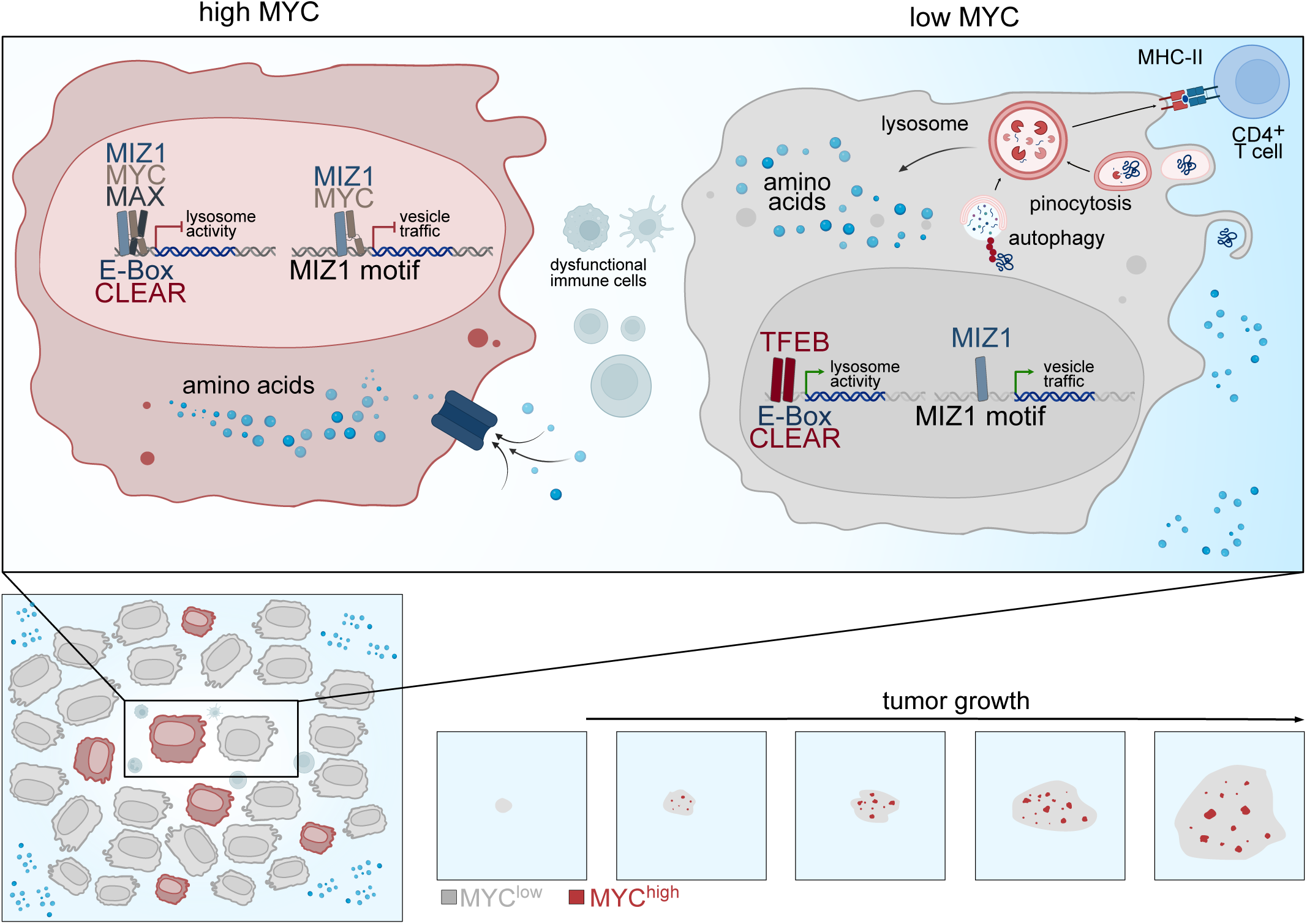

## Introduction

Deregulated MYC expression is a central driver of human tumorigenesis, and genetic models demonstrate that tumors depend on elevated MYC levels throughout cancer progression. Reverting MYC activity in established tumors using conditional alleles or inducible shRNAs leads to rapid regression in multiple solid and hematologic malignancies, including pancreatic ductal adenocarcinomas (PDACs)^1–7^. Expression or delivery of a dominant-negative MYC variant, OmoMYC^3,8^ halts the growth of many tumors, prompting a clinical trial of an OmoMYC peptide^9^.

Elevated MYC expression in PDAC is in part a consequence of the presence of oncogenic KRAS mutations, which drive most PDACs and enhance MYC mRNA abundance and protein stability^10^. MYC is essential for a KRAS-driven oncogenic transcriptional program^11^ and high MYC levels confer resistance to RAS inhibition^12^. Moreover, the *MYC* gene is frequently amplified as extrachromosomal (ec)DNA, leading to high *MYC* copy gene numbers and MYC expression^13^. Intriguingly, both copy number and MYC expression are highly heterogeneous within PDAC tumors, because high MYC levels impose a significant fitness cost on tumor cells^13^, but MYC also promotes PDAC growth by enabling tumors to evade the immune system^14^. Deactivation of MYC in conditional PDAC models leads to infiltration of B and T lymphocytes and natural killer (NK) cells, which contribute to tumor clearance^15,16^. Depletion of endogenous MYC in established tumors from a PDAC model driven by *KRas^G12D^* and *Tp53^R172H^* mutations (“KPC model”)^17^ leads to rapid tumor regression in syngeneic hosts, but MYC is largely dispensable when the same tumors are transplanted into immune-deficient mice^6,18^.

The mechanisms by which MYC drives immune evasion of PDACs are incompletely understood and whether intra-tumoral MYC heterogeneity contributes to this process is unknown. A candidate mediator of MYĆs effects are its well-established roles in reprogramming amino acid metabolism^19^. MYC directly upregulates multiple amino acid transporters, including the glutamine transporter SLC1A5, and induces expression of enzymes that catalyze central steps in amino acid utilization, such as glutamine catabolism^20^. However, PDAC tumors do not rely solely on uptake of free amino acids, since scavenging bulk extracellular protein through macropinocytosis is a dominant nutrient acquisition pathway^21,22^ and autophagy mobilizes intracellular protein stores in parallel^23^. Both macropinocytosis and autophagy are stimulated by KRAS mutations and provide complementary amino-acid sources that compensate for one another when either pathway is impaired^24^.

These pathways converge on lysosomes, which degrade cargo from both macropinocytosis and autophagy and are essential for PDAC growth^25^. Lysosome biogenesis and autophagic flux are controlled by TFEB^26^, MITF^27^, and TFE3^28^, transcription factors that, like MYC/MAX complexes, bind E-box sequences with a core CACGTG motif. In PDAC, these factors are released from regulatory constraints that normally limit their activity, further underscoring the importance of lysosome function^29^. Against this backdrop, it is striking that MYC suppresses TFEB function in acute myeloid leukemia, where TFEB acts as a tumor suppressor^30–32^. Whether MYC attenuates TFEB and lysosome function in PDAC, and by what mechanism, has remained unclear.

A candidate mediator is the POZ/BTB transcription factor MIZ1 (*Zbtb17*), which underlies many of the repressive effects of MYC on transcription^33^. MIZ1 binds its own cognate DNA binding sites and, in the absence of MYC, activates transcription at these loci. Upon complex formation with MYC, MIZ1 forms insoluble complexes that repress transcription from these sites^33^. At high MYC levels, MYC can also recruit MIZ1 to E-box-containing target genes, thereby reducing their transcriptional output^34,35^. Conversely, MIZ1 can recruit MYC to MIZ1 target sites to repress transcription. Direct MIZ1 target genes encode a circumscribed group of proteins involved in membrane transport and vesicle trafficking^5,36^. Consistently, deletion of MIZ1 in neuronal progenitor cells impairs autophagic flux and causes accumulation of polyubiquitylated proteins, indicating that MIZ1 is required for autophagic protein degradation and possibly lysosome function^36^.

Here, single-cell transcriptomic analysis of an orthotopic KPC transplant model was used to define MYC-dependent changes in the tumor microenvironment. These data show that MYC depletes free amino acids in the tumor microenvironment because MYC/MIZ1 complexes repress genes required for lysosome function in tumor cells, thereby limiting lysosomal protein degradation and increasing dependence on uptake of extracellular amino acids. The resulting amino acid scarcity directly impairs the function of multiple immune cell types, causally linking MYC-dependent changes in tumor cell metabolism to MYC-driven immune evasion. We propose that, via this mechanism, whereby high MYC levels compromise lysosome function in a cell-autonomous manner yet confer a non-cell-autonomous advantage, heterogeneity in MYC expression provides a selective advantage to the entire tumor.

## Results

### Depletion of MYC in tumor cells causes CD4^+^ T-cell mediated tumor regression

To profile MYC-dependent changes in the tumor microenvironment (TME) of PDAC, we orthotopically transplanted KPC cells harboring doxycycline-inducible shRNAs targeting MYC into the pancreata of C57BL/6J mice^6,18^ (Supplementary Fig. 1A). Tumors were allowed to establish for 7 days prior to MYC depletion by doxycycline administration (Supplementary Fig. 1B). Consistent with our previous findings, MYC depletion induced robust tumor regression over the subsequent 14 days (Fig. 1A; Supplementary Fig. 1C)^6,18^. To comprehensively profile changes in the TME, we first performed flow cytometry analysis comparing control tumors to those subjected to MYC depletion for seven days (see Supplementary Fig. 1D-F for gating strategy). MYC depletion resulted in increased infiltration of CD8^+^ T cells, CD4^+^ T cells, and B cells (B220^+^), alongside a reduction in monocytes and macrophages (Fig. 1B). To obtain a higher resolution, we next performed single-cell RNA sequencing (scRNA-seq) of sorted immune, tumor, and stromal compartments under the same conditions (Fig. 1C; Supplementary Fig. 2A). This analysis corroborated the flow cytometry results, revealing increased proportions of CD8^+^ T cells, CD4^+^ T cells, and B cells, and decreased abundances of macrophages, monocytes, and granulocytes (including neutrophils and eosinophils) following MYC depletion (Fig. 1D).

**Figure 1:**
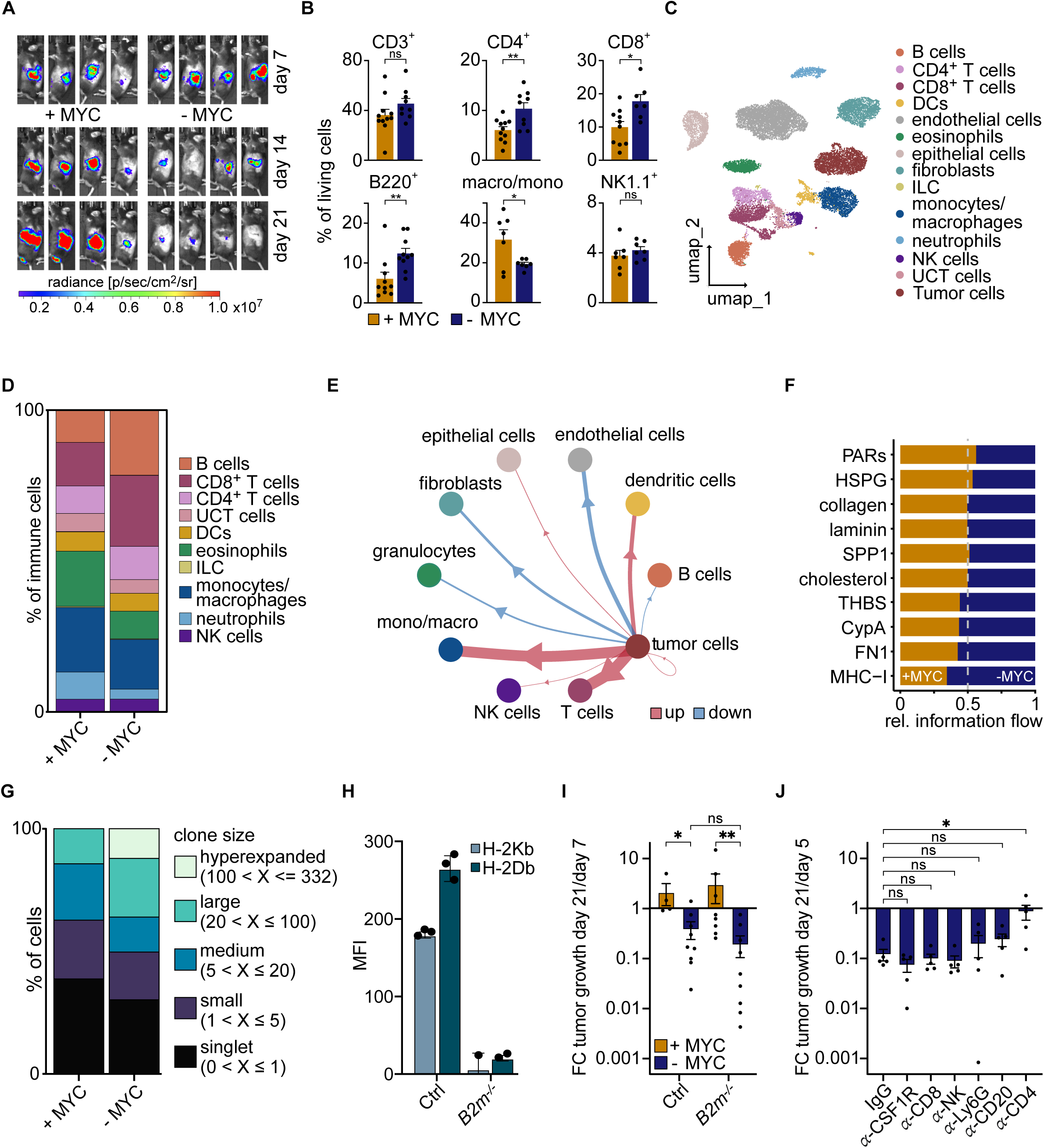
MYC-driven immune evasion is independent of suppression of antigen-presentation via MHC-I. (A) Luciferase imaging of KPC-tumors expressing doxycycline inducible shMYC into C57BL/6J mice at the indicated times after orthotopic transplantation (-MYC) mice received doxycycline from 7 days after transplantation onwards (N = 4). (B) Flow-cytometry analysis of CD3^+^, CD4^+^, CD8^+^, B220^+^cells, macrophages/ monocytes, and NK cells as percentage of live cells. (C) Uniform Manifold Approximation and Projection (UMAP) of scRNA sequencing from indicated cell types from control and MYC-depleted tumors after 7 days of doxycycline treatment (N = 8 mice in each condition, n = 18,614 cells). ILCs: Innate Lymphoid Cells; DC: dendritic cells; UCT: unconventional T cells. (D) Relative distribution of immune cells in the two conditions. (E) Circle plot showing changes in interactions of tumor cells to other cells upon MYC depletion. Clusters with fewer than 50 cells were removed. (F) Relative information flow of tumor-specific intercellular signaling pathways between control and MYC-depleted conditions. (G) Distribution of 5’ TCR single cell sequencing of sorted T cells in control and MYC-depleted tumors categorized based on clonal frequency derived “clone size” as indicated. (N = 8 mice in each condition, n = 4,729 cells). (H) Mean fluorescent intensity (MFI) of KPC cells with CRISPR/Cas9 mediated knockout of *B2m*. Cells were stained for MHC class-I alleles H-2Kb and H-2Db (N = 3; error bars: SD). (I) Fold change (day 21 vs day 7) of tumor growth in +MYC and -MYC tumors, in control and *B2m* knockout conditions (N = 4, 9, 8, 9; Kolmogorov-Smirnov test). (J) Fold change (day 21 vs day 5) of tumor growth of MYC-depleted tumors with antibody depletion of macrophages (α-CSF1R), CD8^+^ T cells, NK cells (α-NK1.1), Neutrophils (α-Ly6G), B cells (α-CD20), and CD4^+^ T cells. Mice were randomized at day 5 and administered doxycycline on day 7 (N=5, Mann-Whitney).

Spatial analyses further refined these observations: immunohistochemistry and spatial transcriptomics demonstrated that B cells preferentially accumulated at the tumor periphery, whereas T cells were distributed throughout the tumor core (Supplementary Fig. 3A-D). To explore how MYC depletion reshapes intercellular communication within the TME, we applied CellChat to infer ligand-receptor interactions across cell populations^37^. MYC depletion enhanced signaling from tumor cells to T cells, dendritic cells, and monocytes/macrophages, while reducing signaling toward fibroblasts, granulocytes, and endothelial cells (Fig. 1E). Among the most prominently enriched pathways in MYC depleted tumors was MHC class I-mediated signaling, consistent with increased antigen presentation by tumor cells (Fig. 1F) and in line with prior reports that MYC suppresses MHC class I expression^18,38–40^. In addition, fibronectin (FN1)- and thrombospondin (THBS)-associated signaling interactions were elevated, consistent with MYC-dependent repression of these pathways^2,16,41^.

These findings suggested that T cells may play a central role in mediating tumor regression following MYC depletion. To assess the functional status and clonality of T cells, we performed 5′ single-cell sequencing to capture T cell receptor repertoires. This revealed clonal expansion of CD8^+^ T cells upon MYC depletion, suggestive of antigen-driven activation (Fig. 1G; Supplementary Fig. 3E). To directly test whether tumor regression depends on MHC class I-mediated antigen presentation, we deleted β2-microglobulin (B2m) in tumor cells (Supplementary Fig. 3F, G), thereby abolishing stable surface expression of MHC class I complexes^25^. Although *B2m* deletion effectively eliminated MHC class I expression (Fig. 1H), it did not impair tumor regression following MYC depletion (Fig. 1I), suggesting that additional MYC-dependent mechanisms regulate anti-tumor immunity. To identify the specific immune cell populations required for this response, we selectively depleted individual immune cell subsets *in vivo* using neutralizing antibodies in tumor-bearing mice followed by depletion of MYC (Supplementary Fig. 3H, I). This approach revealed that CD4^+^ T cells, but not CD8^+^ T cells, NK cells, B cells, macrophages, or neutrophils are essential for tumor regression upon MYC depletion (Fig. 1J).

### Depletion of MYC in tumors activates amino acid metabolism programs in immune cells

To investigate MHC class I-independent mechanisms by which tumor-intrinsic MYC might suppress immune cell function, we analyzed gene expression changes in the immune compartment of the scRNA-seq data. CD8^+^ and CD4^+^ T cells, B cells, and macrophages all responded to MYC depletion in tumor cells with a pronounced increase in the expression of two gene sets encoding proteins involved in amino acid metabolism and protein translation, and, with the exception of B cells, oxidative phosphorylation, indicating a broad reprogramming of amino acid metabolism (Fig. 2A). We hypothesized that increased amino acid availability to immune cells upon MYC depletion in tumor cells could underlie these transcriptional changes. To test this, we performed liquid chromatography-mass spectrometry (LC-MS) of tumor interstitial fluid (TIF) and plasma from tumor-bearing mice at day 14; for plasma, we also included samples from tumor-free mice.

**Figure 2:**
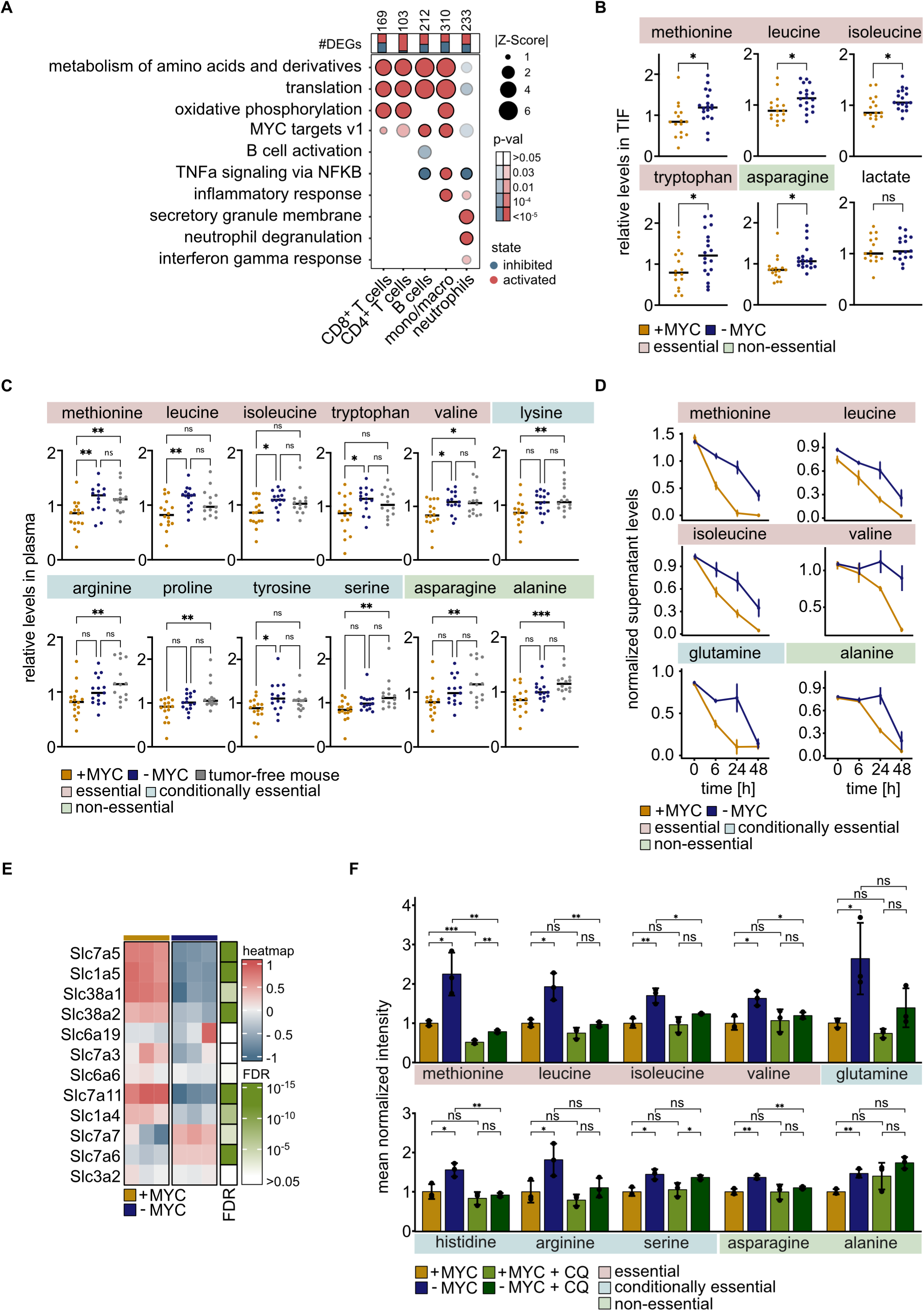
MYC depletion increases local and systemic amino acid levels. (A) Over-representation analysis of differentially expressed genes in the indicated immune cell compartments after MYC depletion (-MYC) in tumors for 7 days, compared to control (+MYC). State indicates predicted activation or inhibition of a pathway in MYC depleted tumors. Top top panel shows number of differentially expressed genes (FDR < 5%) with bars depicting the proportion of up- and downregulated genes. (B) LC-MS analysis of interstitial fluid from tumors before and after MYC depletion (N = 17,18; unpaired, two-sided t-test). (C) LC-MS analysis of plasma from tumor-bearing mice (+/- MYC) and healthy, age and sex-matched mice (N = 16, 15, 14; ordinary, one-way ANOVA). (D) LC-MS analysis of supernatant of KPC cells cultivated in human-plasma-like medium (HPLM). Cells were grown to confluence; medium was refreshed, and supernatant was sampled at timepoint 0 h, 6 h, 24 h, and 48 h. Values are normalized to levels in the medium (N = 3, error bars: SD). (E) Heatmap of RNA expression of amino acid transporters in KPC cells after shRNA mediated depletion of MYC for 48 h (N = 3). FDR from differential gene expression analysis annotated on the right panel. (F) Bar plot shows LC-MS analysis of intracellular metabolites in KPC cells. Values are normalized to the + MYC condition. Cells were treated with doxycycline to deplete MYC for 48 h and treated with 25 µM chloroquine (CQ) for the last 4 h. (N = 3; error bars: SD; unpaired t-test).

Depletion of MYC in tumor cells did not significantly change the weight of the tumors (Supplementary Fig. 1C) but led to a significant increase in the levels of methionine, leucine, isoleucine, tryptophan, and asparagine in the TIF (Fig. 2B), whereas levels of other amino acids did not change significantly (Supplementary Table 1). These effects were specific, as MYC depletion did not alter lactate, aldohexose, or other measured metabolites (Fig. 2B, Supplementary Fig. 4A, Supplementary Table 1). Surprisingly, MYC-dependent changes in amino acid concentrations were not restricted to the tumor microenvironment and were also detectable in plasma (Fig. 2C). Relative to tumor-free mice, tumor-bearing mice displayed significantly reduced levels of multiple essential, conditionally essential, and non-essential amino acids in the plasma, while depletion of MYC in established tumors partly or completely restored these plasma amino acid concentrations toward normal (Fig. 2C, Supplementary Table 2).

To demonstrate that these effects directly arise from MYC-dependent changes in amino acid uptake and/or metabolism by tumor cells and exclude confounding effects of different tumor sizes *in vivo*, we performed LC-MS analysis of culture supernatants from KPC tumor cells grown to confluence to equalize cell number across conditions. This experiment showed that KPC cells rapidly deplete the medium of multiple amino acids in a MYC-dependent manner (Fig. 2D). We confirmed previous observations that multiple amino acid transporters are expressed in a MYC-dependent manner (Fig. 2E). Despite the decrease in extracellular amino acid uptake upon MYC depletion, intracellular concentrations of multiple amino acids increased (Fig. 2F), raising the possibility that high MYC activity limits the availability of endogenous amino acid pools in KPC cells. Strikingly, the replenishment of intracellular amino acid pools after MYC depletion was abrogated by brief incubation with chloroquine (CQ), a drug that impairs lysosomal acidification and function (Fig. 2F; Supplementary Table 3). This finding suggests that endogenous MYC attenuates lysosomal function and the utilization of lysosome-derived amino acids in KPC cells, prompting us to further probe the underlying mechanisms.

### Complex formation of MYC with MIZ1 attenuates lysosomal protein degradation

RNA-sequencing revealed that MYC depletion in KPC cells both down- and upregulated large groups of genes (Fig. 3A). Gene that were downregulated upon MYC depletion were enriched in canonical MYC targets and cell cycle regulators, consistent with previous work and arguing that MYC activates these genes (Fig. 3B)^5,7^. In contrast, genes upregulated upon MYC depletion were highly enriched for lysosomal and autophagy-related functions, suggesting that MYC transcriptionally suppresses these processes (Fig. 3A, B). Consistent with this, MYC-depleted cells displayed increased LC3 lipidation, particularly in the presence of chloroquine, a pattern consistent with increased autophagic flux (Fig. 3C). Amino acid starvation further enhanced LC3-II accumulation in MYC-depleted or MYC-deleted cells, whereas cells with unperturbed MYC failed to mount a comparable response (Supplementary Fig. 4B).

**Figure 3:**
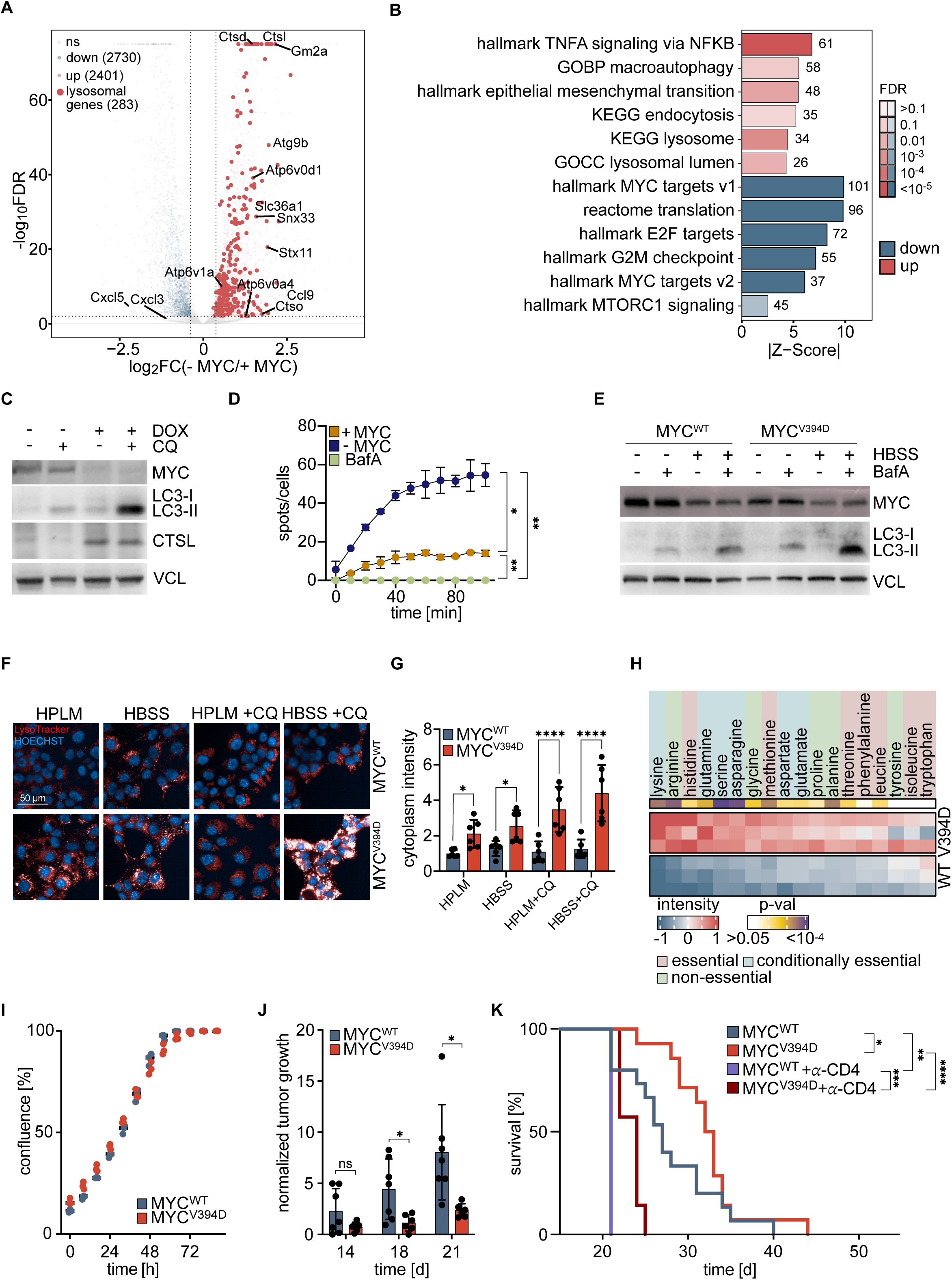
MYC/MIZ1 interaction inhibits lysosome function and sustains tumor growth. (A) Volcano plot of differentially expressed genes upon MYC depletion (FDR < 1%; FC>1.3). Genes associated with lysosomal functions ar highlighted and exemplary genes are labeled. (B) Over-representation analysis of differentially expressed genes in A. State indicates predicted activation or inhibition of a pathway. Numbers indicate number of genes in respective pathways. (C) Immunoblot of MYC, LC3, CTSL in +MYC and -MYC KPC cells treated with doxycycline (1 µg/mL, 48 h) to deplete endogenous MYC or EtOH as control. Indicated samples were also treated with chloroquine (4h) before precipitation. Vinculin is loading control (N = 2). (D) Number of lysosomes per cell from live-cell imaging stained with DQ-Red. Cells treated with bafilomycin A (BafA, 100 nM, 4 h) were used as control (N = 2, t-test). (E) Immunoblot of MYC and LC3 in KPC cells with depletion of endogenous MYC and expression of doxycycline inducible MYC^WT^ or MYC^V394VD^ cultivated in normal medium (HPLM) or under starvation in Hanks’ Balanced Salt Solution (HBSS). Indicated samples were also treated with BafA. Vinculin is loading control. (F) Lysotracker-staining of KPC cells expressing MYC^WT^ or MYC^V394VD^ and depletion of endogenous MYC cultivated in HPLM or HBSS. Scale bar indicates 50 µM. Nuclei were visualized using Hoechst. (G) Quantification of (F) (N = 6; error bars: SD; Welch’s t-test). (H) LC-MS analysis of water-soluble metabolites in cells expressing MYC^WT^ or MYC^V394D^. Heatmap shows relative normalized peak areas, p-values generated from unpaired t-tests on normalized peak areas. (I) Growth curve of KPC cells expressing doxycycline-inducible shRNA targeting endogenous MYC and doxycycline-inducible expression of MYC^WT^ and MYC^V394D^ (N = 3). (J) Normalized tumor growth of KPC tumors expressing MYC^WT^ or MYC^V394D^. (K) Kaplan-Meier plot of tumor-bearing mice with tumor cells expressing MYC^V394D^. Tumors engrafted for 5 days and were then randomized and treated with IgG or α-CD4 antibody. Mice received doxycycline from day 7 on (N = 15, 14, 7, 5 mice, Logrank (Mantel-Cox)).

Depletion of MYC also increased mRNA and protein levels of the lysosomal protease cathepsin L (CTSL) (Fig. 3A, C). To directly assess lysosomal protease activity, we used DQ-Red, a self-quenching fluorogenic substrate that becomes fluorescent upon proteolytic cleavage by lysosomal cathepsins^42^. Live-cell imaging revealed that MYC depletion increased the number of DQ-Red positive lysosomes compared to control cells with unperturbed MYC, and this increase was blocked by treatment with bafilomycin A (BafA), an inhibitor of vacuolar ATPases and lysosome acidification (Fig. 3D). Furthermore, MYC depletion enhanced uptake of fluorophore-coupled BSA, and this increase was abrogated by the macropinocytosis inhibitor 5-(N-ethyl-N-isopropyl)amiloride (EIPA) (Supplementary Fig. 4C). Together, these data show that endogenous MYC limits autophagy, lysosomal protease activity, and macropinocytosis in KPC cells.

MYC can suppress transcription through association with the POZ/BTB domain transcription factor MIZ1 (see Introduction). To test whether MYC/MIZ1 complexes repress lysosome function, we performed RNA-sequencing on KPC cells expressing either wild-type MYC (MYC^WT^) or MYC^V394D^, a point mutant that retains its MAX interaction but is impaired in binding to MIZ1^33^. In these cells, endogenous MYC was depleted using doxycycline-inducible shRNAs, and MYC^WT^ or MYC^V394D^ was expressed at comparable levels (5) (Fig. 3E). MYC^V394D^ cells upregulated multiple lysosomal genes relative to MYC^WT^ cells (Supplementary Fig. 4D, E). In line with this, MYC^V394D^-expressing cells displayed enhanced LC3 lipidation upon bafilomycin A treatment that further increased in amino acid-free medium, consistent with increased autophagic flux (Fig. 3E). LysoTracker staining showed that MYC^WT^-expressing cells harbored few cytosolic lysosomes which did not increase upon amino acid starvation or chloroquine treatment, whereas MYC^V394D^-expressing cells showed strongly elevated staining under these conditions (Fig. 3F, G). Reduced lysosome function in MYC^WT^ cells also correlated with markedly decreased intracellular amino acid levels compared with MYC^V394D^ cells (Fig. 3H). These data further supported our hypothesis that MYC/MIZ1 complexes attenuate lysosome function and limit amino acid availability in KPC cells.

In culture, KPC cells expressing MYC^V394D^ grew at the same rate as cells expressing MYC^WT^, underlining that MYC^V394D^ is capable to activate MYC-dependent proliferation programs (Fig. 3I). In contrast, MYC^V394D^ tumors grew slower than MYC^WT^ tumors upon orthotopic transplantation into C57BL/6J mice, and mice bearing MYC^V394D^ tumors survived significantly longer than those bearing MYC^WT^ tumors (Fig. 3J, K). Depletion of CD4^+^ T cells shortened survival in both groups and reduced the average survival difference between MYC^WT^ and MYC^V394D^ bearing mice from about 6 days to 3 days (Fig. 3K). Flow cytometric analysis of *in vivo* tumor cells revealed robust MHC class II expression on their surface, suggesting that direct CD4^+^ T cell-mediated cytotoxicity against tumor cells is at least feasible (Supplementary Fig. 4F, G). We concluded that MYC/MIZ1 complexes are dispensable for KPC cell proliferation in culture, but promote tumor growth *in vivo*, likely by suppressing CD4^+^ T cell-mediated antitumor immunity.

### MYC/MIZ1 complexes repress lysosome function via both TFEB and MIZ1 target sites

Analysis of available ChIP-sequencing data^34^ identified 7,051 promoters that are occupied by both MYC and MIZ1 (Fig. 4A). Comparison of these with published CHIP enrichment analysis (ChEA) and RNA-sequencing datasets showed that 1,412 MYC-repressed genes have promoters co-occupied by MYC and MIZ1 (Fig. 4B). Of these, 173 are known direct targets of the TFEB/TFE3/MITF network, a central regulator of lysosome biogenesis^26,43^. A further 274 MYC-repressed genes were identified as MIZ1-dependent by RNA-seq upon MIZ1 knockdown^16^, with only 25 genes overlapping between the TFEB/TFE3/MITF and MIZ1-dependent sets (Fig. 4B). Functional annotation showed that both classes are highly enriched for lysosome-related genes, suggesting that MYC/MIZ1 complexes suppress lysosomal activity through both TFEB-responsive and MIZ1-responsive target sites (Fig. 4C). Motif analysis and genome browser inspection further indicated that promoters of lysosome-associated genes co-bound by MYC and MIZ1 contain either the CLEAR/E-box motif recognized by TFEB/TFE3/MITF or a MIZ1 binding motif (Fig. 4D, E).

**Figure 4:**
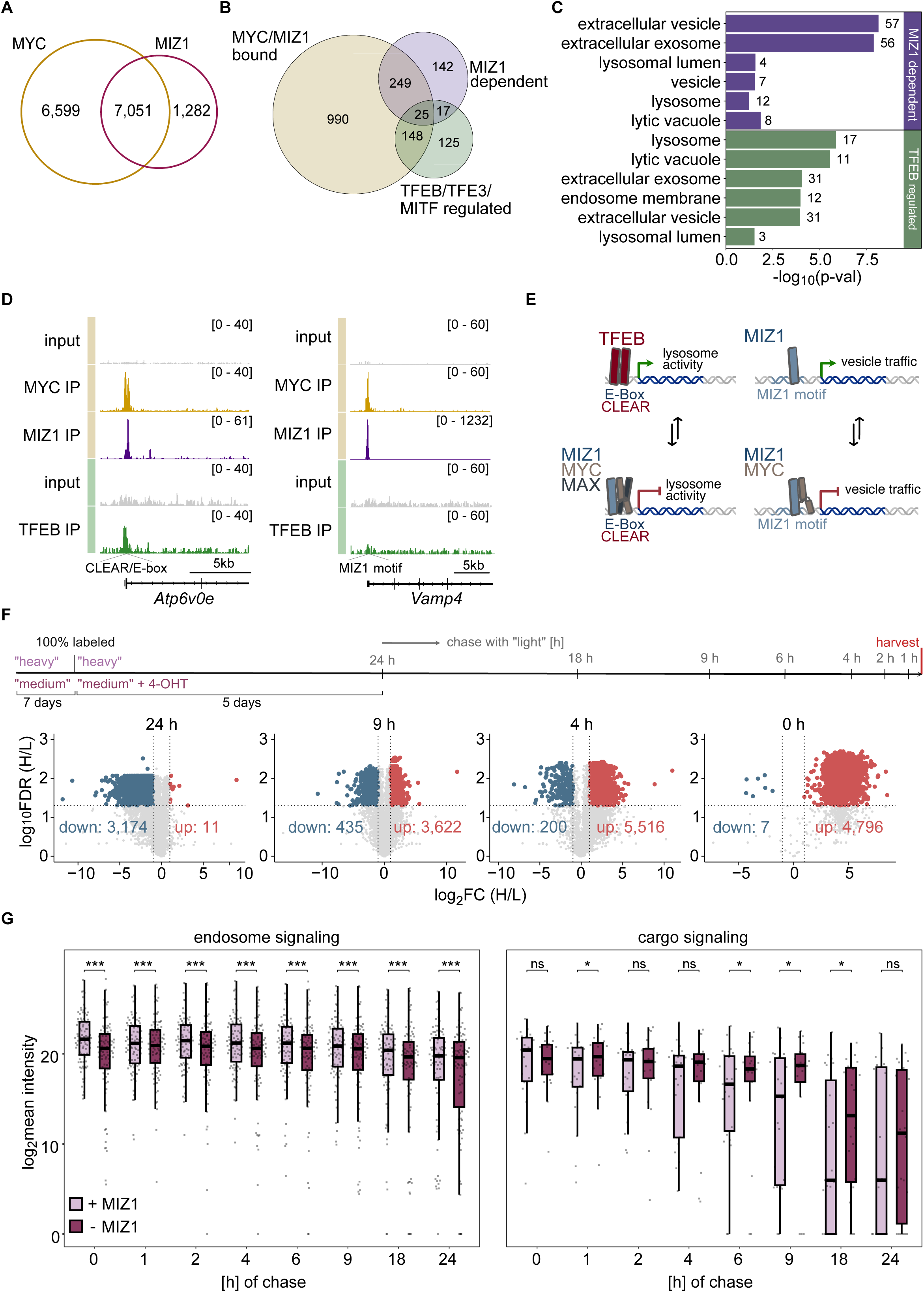
MYC/MIZ1 complex represses classes of genes involved in lysosomal degradation and endosomal sorting/trafficking. (A) Venn diagram showing MYC and MIZ1 bound promoters. (B) Venn diagram with genes upregulated upon MYC depletion that are MYC/MIZ1 bound, MIZ1 dependent, and/or TFEB/TFE3/MITF regulated. (C) Over-representation analysis of MIZ1 dependent (249 genes) and TFEB regulated genes (148 genes) from (B) that are MYC/MIZ1 bound but mutually exclusive of each other. (D) Exemplary browser tracks of a TFEB-regulated gene and a MIZ1-regulated gene, both bound and repressed by MYC/MIZ1. (E) Scheme showing two mechanisms by which MYC/MIZ1 could repress genes involved in the broader umbrella of lysosome function. (F) Schematic for SILAC proteomics of control and MIZ1 knocked out cells (+ 4-OHT) labeled with “heavy” and “medium” animo acids respectively and chased with “light” over different timepoints. Exemplary volcano plots from different timepoints tracking labeled vs unlabeled proteins in the control cells serve as quality control. (G) Box plots showing proteins corresponding to endosomal function and cargo/signaling proteins in control and MIZ1-knocked out cells across indicated timepoints. Wilcoxon rank sum test was used to compute statistical significance between - MIZ1 vs MIZ1 (N = 3).

To test whether MIZ1-activated genes are required for lysosomal protein degradation in KPC cells, we generated a conditional MIZ1 (*Zbtb17*) allele in which exons 7-9 of *Zbtb17* are flanked by loxP sites, crossed this allele into KPC mice expressing CreER, and established cell lines from tumors that developed in the absence of 4-hydroxytamoxifen (4-OHT) (Supplementary Fig. 5A). PCR and RNA-seq tracks confirmed efficient deletion of exons 7–9 upon 4-OHT treatment (Supplementary Fig. 5B, C). To globally measure the effect of MIZ1 on protein stability in an unbiased manner, we performed an arginine/lysine-based SILAC pulse-chase experiment in control and 4-OHT-treated cells. Cells were metabolically labeled with “medium” (control) or “heavy” (4-OHT) isotopic Arg/Lys and subsequently chased in medium containing unlabeled (“light”) amino acids for 0-24 h (Fig. 4F). Volcano plots confirmed the expected shift from labeled to unlabeled peptides between 0 h and 24 h, consistent with global protein turnover (Fig. 4F). To dissect compartment-specific effects, we then compared labeled protein abundance at each chase time point and carried out over-representation analysis of differentially abundant labeled proteins. Endosome-related terms were significantly underrepresented at 0 h in MIZ1-deleted cells compared to control but appeared among proteins that were both up and down at later chase times (e.g., after 6 h) (Supplementary Fig. 6D), prompting closer examination of individual proteins in these pathways. Detailed functional annotation revealed that this bidirectional enrichment reflected two opposing programs: deletion of MIZ1 reduces endosomal signaling and core machinery proteins already before the start of the chase and remained significantly decreased throughout, whereas endosomal cargo and signaling substrates of this machinery were more stable upon MIZ1 deletion, indicating impaired degradation (Supplementary Table 4, Fig. 4G). Together with the genomic data, these findings support a model in which MYC attenuates lysosomal function and lysosomal protein degradation by repressing TFEB-network and MIZ1-dependent target genes pivotal for lysosome function.

### Replacement of mouse by human Na^+^/K^+^-ATPase sensitizes KPC cells to cardiac glycosides

Since MYC has pleiotropic effects on tumor cells, we sought an orthogonal strategy to validate whether amino acid depletion in the tumor microenvironment mediates immune evasion downstream of MYC. To accomplish this, we engineered a system that can selectively and acutely block amino acid uptake in tumor cells. Because amino acid uptake depends on the Na^+^ gradient, we inhibited Na^+^/K^+^-ATPase activity using cardiac glycosides such as digitoxin, digoxin, and cymarin (Fig. 5A). These cardiac glycosides have comparable K_d_ values for human Na^+^/K^+^-ATPase and were used interchangeably in our experiments. To specifically inhibit amino acid import into tumor cells, we exploited the observation that the human Na^+^/K^+^- ATPase is sensitive to cardiac glycosides, while the murine protein harbors two amino acid changes that reduce the affinity to cardiac glycosides ∼1,000fold^44^. We therefore generated KPC^huATP1A1^ cells that expressed human ATP1A1, encoding the α1 subunit, and used CRISPR/Cas9 to delete the endogenous murine *Atp1a1* allele, rendering them sensitive to cardiac glycosides (Supplementary Fig. 6A). Orthotopic transplantation of KPC^huATP1A1^ cells into mice thus created a hybrid system that permits selective inhibition of amino acid import into tumor cells upon treatment, while sparing host immune cells (Fig. 5A).

**Figure 5:**
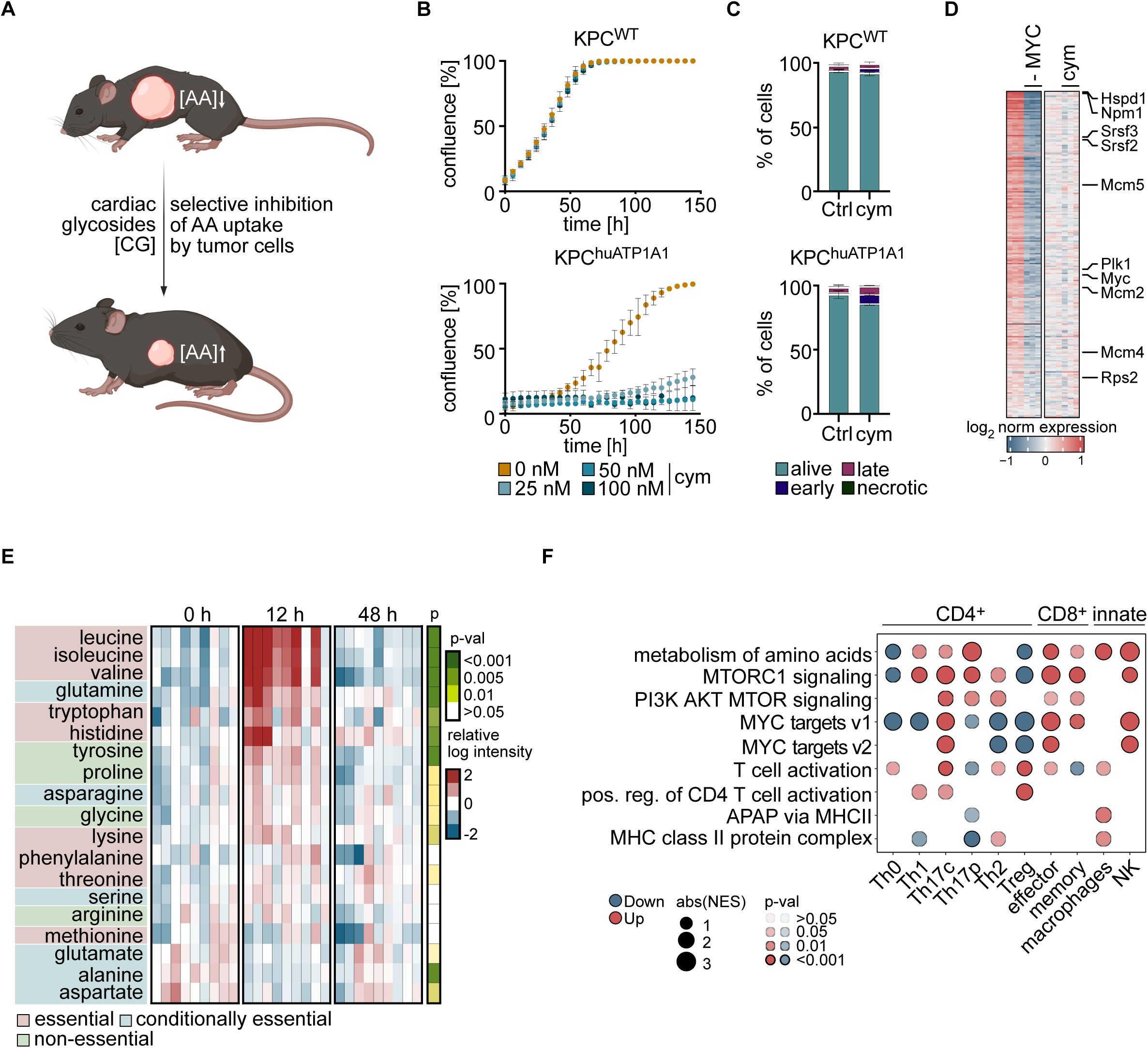
Amino acid levels control immune cell function in culture. (A) Diagram illustrating the generation of KPC cells with CRISPR/Cas9-mediated deletion of endogenous murine *Atp1a* that express human ATP1A1 (KPC^huATP1A1^). Addition of cardiac glycosides inhibits import of amino acids into tumor cells without affecting host cell. (B) Growth of KPC^WT^ and KPC^huATP1A1^ in the presence of the indicated cymarin concentrations (N = 3; error bars: SD). (C) Annexin-PI flow cytometry analysis of KPC^WT^ and KPC^huATP1A1^ cells treated for 48 h with 100 nM of cymarin (N = 3; error bars: SD). (D) Heatmap of genes of MYC targets V1 from RNA sequencing of tumor cells with depletion of MYC using shRNAs or treatment of KPC^huATP1A1^ cells with 100 nM cymarin for 24 h (N = 3). (E) Heatmap of amino acid levels in the interstitial fluid of tumors engrafted from KPC^huATP1A1^ cells upon treatment with digitoxin (2 mg/kg body weight) for the indicated time (N = 9). (F) Over-representation analysis of mRNA sequencing of different immune cells isolated and cultured *ex vivo* and supplemented with amino acids mimicking their concentrations in the TIF observed in (D) at 12 h of CG treatment.

In culture, treatment of human PDAC cells (BxPC-3 and PaTu 8988T) with cymarin markedly reduced steady-state levels of amino acids and derived metabolites, including purines and pyrimidines, while wild-type KPC cells showed no significant changes (Supplementary Fig. 6B). This metabolic depletion correlated with reduced proliferation of PaTu 8988T and Panc1 cells, as well as KPC^huATP1A1^, but not KPC^WT^ cells (Supplementary Fig. 6C, Fig. 5B). The delay in proliferation similarly affected all phases of the cell cycle (PaTu 8988T, Panc1 cells: Supplementary Fig. 6D; KPC^huATP1A1^, KPC^WT^: Supplementary Fig. 6E). In contrast, cardiac glycosides did not induce apoptosis of murine cells (Fig. 5C). RNA-sequencing and immunoblotting showed that digitoxin did not cause major changes in canonical MYC target gene expression (Fig. 5D) or activation of TBK1 (Supplementary Fig. 6F). To directly test effects on amino acid uptake, we quantified metabolites in culture supernatants of KPC^huATP1A1^ cells by LC-MS. Concentrations of multiple amino acids, particularly methionine, alanine, leucine, isoleucine, and valine, decreased in the supernatant of control cells but declined much less in cymarin-treated cells, indicating reduced uptake (Supplementary Fig. 7A).

We next orthotopically transplanted KPC^huATP1A1^ cells into C57BL/6J mice and treated animals with digitoxin one week after engraftment. Digitoxin caused a transient increase in several amino acids, including leucine, isoleucine, valine, and glutamine, in TIF, whereas lactate and carbohydrate levels remained largely unchanged (Fig. 5E, Supplementary Fig. 7B). Amino acid concentrations rose significantly at 12 h but returned to baseline by 48 h (Fig. 5E). To determine whether this metabolic cue was sufficient to reprogram immune cells, we cultured a panel of purified immune cells in HPLM and treated them with an amino acid cocktail recapitulating the TIF composition at 12 h after digitoxin. RNA-sequencing revealed that the cocktail regulated gene sets such as amino acid metabolism, translation, and MYC targets, consistent with a direct transcriptional response to increased amino acid availability and subsequent acquisition of activation-associated programs (Fig. 5F). However, this response was cell-type specific rather than global: effector and inflammatory populations (Th1, Th17c, CD8^+^ effector and memory T cells, NK cells and macrophages) upregulated mTORC1 signaling and MYC targets, whereas naive (Th0) and regulatory (Treg) T cells showed the opposite response, downregulating the same pathways. Macrophages additionally upregulated antigen-presentation and MHC class II programs (Fig. 5F). This showed that the increasing amino acid concentrations in the tumor microenvironment likely did not serve as a global, uniform stimulant, but targeted differentiating and effector like populations while sparing the rest.

### Transient inhibition of amino acid import causes tumor eradication and long-term survival

Treatment of C57BL/6J mice bearing KPC^huATP1A1^ tumors with digitoxin led to marked tumor regression, as quantified by tumor weight (Fig. 6A). Inspection of histology showed near-complete disintegration of tumor tissue after two days of treatment (Fig. 6B). Multicolor immunofluorescence revealed migration of T cells from the tumor periphery into the tumor core and an increase in F4/80^+^ and CD169^+^ macrophages within the tumor, while the number of cytokeratin-positive tumor cells declined rapidly (Fig. 6C).

**Figure 6:**
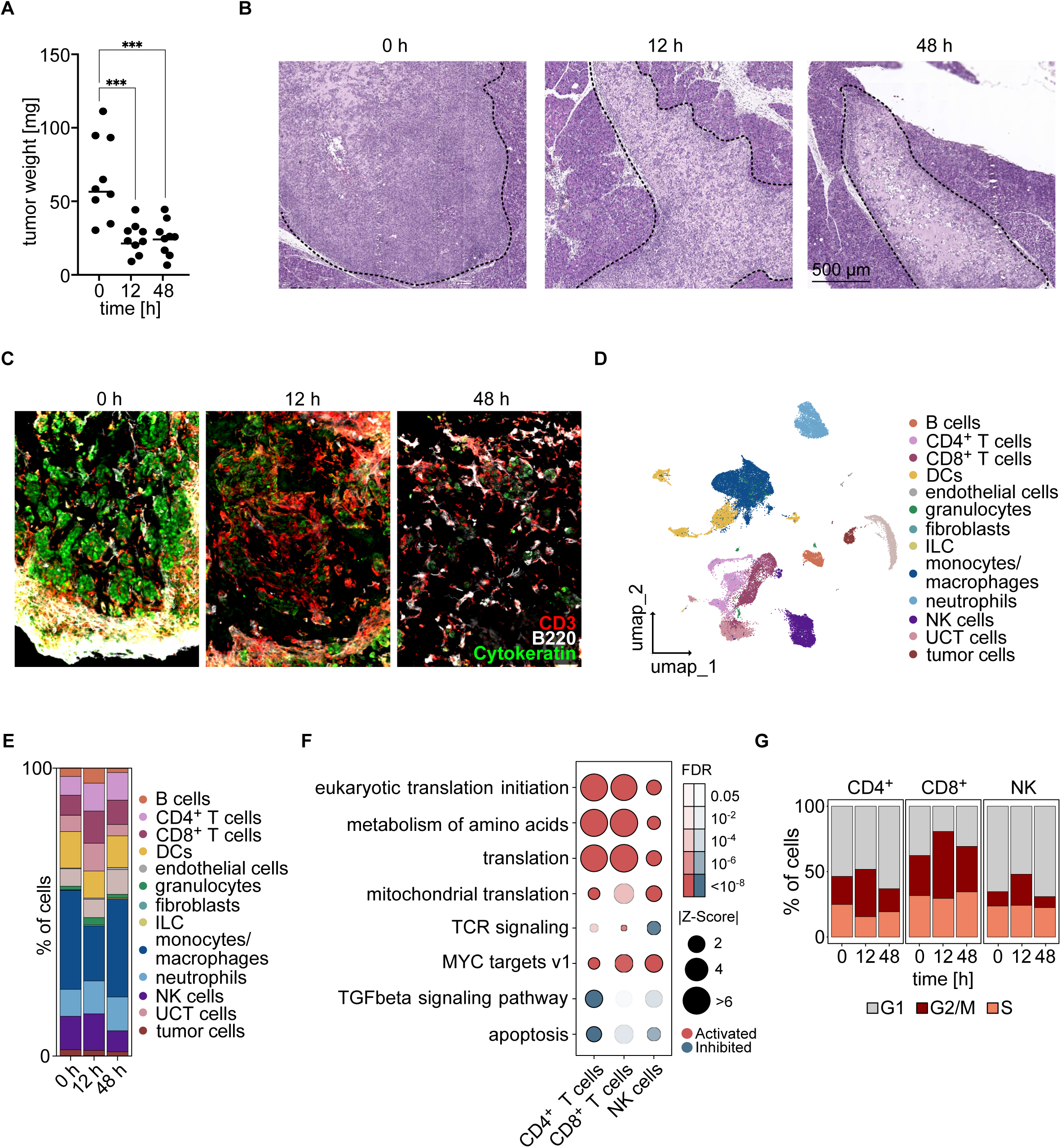
Inhibition of amino acid import into tumors restores immune surveillance. (A) Weight of tumors from KPC^huATP1A1^ cells. Mice were treated with digitoxin for the indicated times after engraftment for seven days (N = 9; ordinary one-way ANOVA). (B) Hematoxylin and eosin (H&E) staining of tumors engrafted from KPC^huATP1A1^ cells. Mice were treated with digitoxin for the indicated time. Tumor area is highlighted with a dashed line (N = 9). (C) Immunofluorescence staining of tumors engrafted from KPC^huATP1A1^ cells. Mice were treated with digitoxin for the indicated time (N = 3). (D) UMAP of scRNA sequencing after engraftment of KPC^huATP1A1^ cells. (E) Relative frequency of the indicated cell types before and after digitoxin treatment (2 batches, N = 4 mice each, n = 38,608 cells). (F) Over-representation analysis of differentially expressed genes in the indicated cell types in response to digitoxin at 12 h vs 0 h. (G) Cell cycle distribution of the indicated cell types, as inferred from the scRNA sequencing.

scRNA-seq of tumor-infiltrating lymphocytes and epithelial cells showed an early increase in the relative abundance of T cells and a concomitant decrease in macrophages within 12 h of digitoxin treatment. These changes were transient and reverted to control values by 48 h, coinciding with the rapid reduction in tumor mass (Fig. 6A, D, E). Importantly, gene-expression profiles across multiple immune cell populations revealed increased expression of gene sets involved in amino acid metabolism, protein translation, and oxidative phosphorylation, closely recapitulating the transcriptional changes observed in lymphocytes after MYC depletion in tumor cells (Fig. 6F). In addition, both CD4^+^ and CD8^+^ T cells and NK cells displayed a higher fraction of cells in S or G2/M phase, indicating increased proliferation (Fig. 6G). Notably, shifts in immune-cell composition and proliferation status also normalized by 48 h, consistent with the near-complete disappearance of tumor tissue at this time point (Fig. 6 A, B, E).

To examine long-term consequences of selectively inhibiting amino acid import in tumor cells, we then transplanted KPC^huATP1A1^ cells into immunocompromised NRG mice. After seven days of tumor growth, we initiated digitoxin treatment and monitored tumor burden weekly by bioluminescence imaging (Fig. 7A). In NRG mice, digitoxin only modestly delayed tumor growth (Fig. 7A, Supplementary Fig. 7C). In sharp contrast, digitoxin induced rapid and profound tumor regression in syngeneic, immune-competent C57BL/6J mice; after seven days of treatment, tumors were no longer detectable (Fig. 7B, Supplementary Fig. 7C). Prolonged treatment for three additional weeks provided only a marginal survival benefit in NRG mice, whereas digitoxin-treated C57BL/6J mice experienced no tumor recurrence and all animals remained tumor-free up to 365 days (Fig. 7C). Depletion of specific immune populations in C57BL/6J mice revealed that loss of CD4^+^ T cells, and to a lesser extent B cells, abrogated or impaired digitoxin-induced tumor eradication, whereas depletion of CD8^+^ T cells or NK cells had minimal impact (Fig. 7D, Supplementary Fig. 7D, E). Collectively, the data show that transient inhibition of amino acid import into tumor cells causes CD4^+^ T cell-mediated eradication of murine PDACs and enables long-term survival of PDAC-bearing mice.

**Figure 7:**
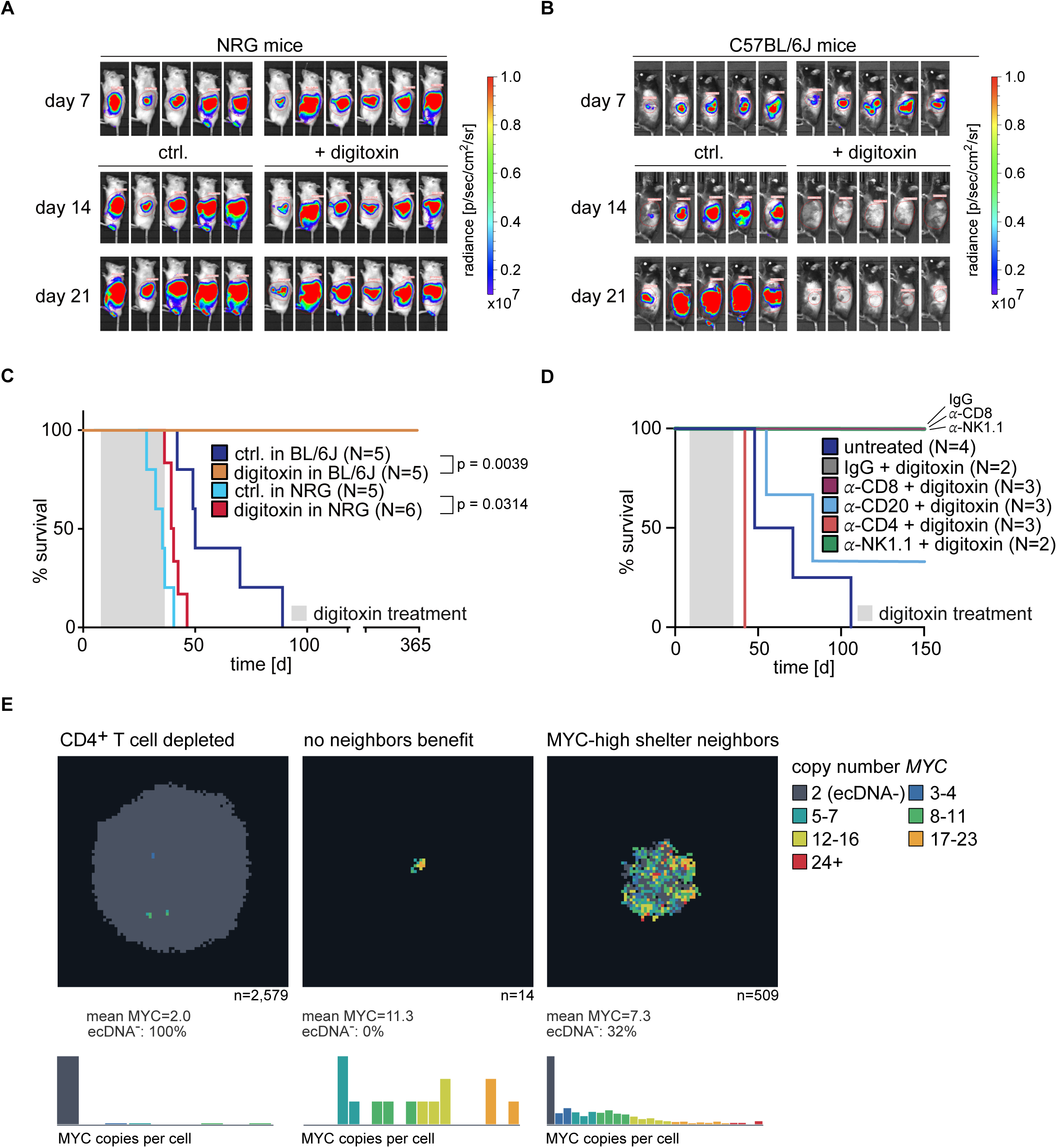
Selective inhibition of amino acid transport induces immune-dependent regression of PDAC tumors. (A) Luciferase imaging of KPC^huATP1A1^ cell-derived tumors in NRG mice. Mice were randomized on day 7 and treated with digitoxin or vehicle. Tumor size was monitored on day 14 and 21 (N = 5, 6). (B) Luciferase imaging of KPC^huATP1A1^ cell-derived tumors in C57BL/6J mice. Mice were treated as in (A) (N = 5, 5). (C) Kaplan-Meier plot of tumor-bearing mice with KPC^huATP1A1^-cell-derived tumors in NRG mice and C57BL/6J mice. Mice were treated with digitoxin for 28 days (Gehan-Breslow-Wilcoxon test). (D) Kaplan-Meier plot of tumor-bearing mice with KPC^huATP1A1^-cell-derived tumors (Gehan-Breslow-Wilcoxon test). Tumors were engrafted in C57BL/6J mice. Mice were treated with digitoxin and IgG, α-CD8, α-CD4, α-CD20, and α-NK1.1 antibodies. (E) Agentic AI-aided simulation, illustrating the proposed mechanism where a growing tumor in which ecDNA-borne MYC is randomly segregated at each division and imposes a fitness cost. The model is intended as a schematic illustration of the hypothesis; protection parameters are constructed rather than measured.

### Combined inhibition of lysosome function and amino acid depletion selects for heterogeneous MYC expression

Lysosome function is central to KRAS-driven nutrient acquisition in PDAC, so its inhibition by MYC is expected to impose a fitness burden on tumor cells *in vivo*. We therefore hypothesized that a mechanism in which MYC imposes a cell-autonomous fitness cost yet provides non-cell-autonomous protection from immune killing would select for heterogeneous MYC expression in PDAC. Since MYC is frequently carried on ecDNA^13^, we explored this by implementing a minimal spatial agent-based model in which ecDNA-borne *MYC* is randomly segregated at division, incurs a dosage-dependent cost, and confers protection from immune-mediated killing to MYC-high cells and their immediate neighbors (Supplementary Fig. 7F). Supplementary Data 1 provides an HTML interface that allows visualizing tumor growth and *MYC* copy number across a wide range of parameters. The model predicts the growth of large tumors with low *MYC* copy numbers in the absence of immune cells, whereas immune cells without MYC-mediated bystander protection potently suppress tumor growth (Fig. 7E). By contrast, when MYC-high cells protect their neighbors, large tumors grow which select for highly variable *MYC* levels (Fig. 7E). Thus, this mechanism provides a formal explanation for the MYC expression heterogeneity observed in human PDACs.

## Discussion

MYC-driven tumors exhibit characteristic hallmarks across multiple entities^45^. Here, we show that two of these hallmarks, the ability of MYC to drive immune escape and to enhance uptake of free amino acids, are mechanistically linked, as MYC-dependent amino acid depletion in the tumor microenvironment directly compromises immune cell function.

Tumor cells import free amino acids via solute carrier proteins, and MYC stimulates uptake and utilization of amino acids by inducing the expression of transporter genes such as SLC1A5^20^. In addition, PDAC tumors use lysosomal degradation of proteins derived from autophagy and macropinocytosis as redundant routes to acquire amino acids^24^. We found that MYC attenuates lysosome function in KPC tumors through MIZ1, a partner that mediates many repressive effects of MYC on transcription^34,46^, including repressive effects on NF-kB and IRF-dependent gene expression during PDAC development^16,35,47^. MYC/MIZ1 complexes repress genes critical for autophagy and lysosome function in two ways. First, MYC recruits MIZ1 to E-box target sites, many of which are bound and activated by TFEB, a master regulator of lysosome biogenesis. Second, MIZ1 recruits MYC to repress MIZ1 target genes involved in vesicular transport that are required for degradation of lysosomal cargo proteins. These findings align with reports that MYC attenuates TFEB function in acute myeloid leukemia^30–32^ and show that high MYC levels suppress lysosomal protein degradation in PDAC tumors.

As consequence of both effects, KPC tumors broadly deplete amino acids from the microenvironment in a MYC-dependent manner, and immune cells respond to MYC depletion in tumor cells by upregulating amino acid-metabolism programs. These findings are surprising, since lysosome function, autophagy and macropinocytosis are stimulated by KRAS and are critical for nutrient acquisition of KRAS-driven PDAC tumors^24^. PDAC cells also divert MHC class I molecules to lysosomal degradation, directly linking lysosomes to immune evasion^25^. We therefore used an orthogonal genetic system to firmly establish that depletion of amino acids indeed is critical for immune evasion. To do so, we exploited the differential sensitivity of mouse and human Na^+^/K^+^-ATPase to cardiac glycosides (CG), generating hybrid mice in which CG treatment selectively inhibits amino acid uptake by tumor cells while sparing host cells. In these mice, CG treatment rapidly increased amino acid levels in the tumor microenvironment, consistent with reduced tumor cell uptake—without significantly affecting expression of MYC target genes. Strikingly, CG-mediated inhibition of amino acid import into tumors transplanted into immunocompetent mice caused complete tumor regression and long-term survival, whereas cardiac glycosides had virtually no effect on KPC tumor growth in immune-deficient mice, likely because residual import sufficed to sustain growth in the absence of immune pressure. CD4^+^ T cells were essential for CG-mediated tumor eradication, and the magnitude of amino acid increase observed *in vivo* was sufficient to activate CD4^+^ T cells in culture. These data establish that counteracting MYC-dependent amino acid depletion in the tumor microenvironment restores immune eradication of tumor cells in this model.

Collectively, these data show that MYC and mutant KRAS drive opposing programs of amino-acid acquisition, each conferring a distinct selective advantage to PDAC cells. One model to explain these findings posits that MYC-driven depletion of extracellular amino acids acts in a non-cell-autonomous manner, whereas suppression of lysosome function impairs amino-acid acquisition from protein degradation in a cell-autonomous manner. MYC expression in human PDACs is highly dynamic and heterogeneous at the single-cell level, particularly in tumors that carry MYC on ecDNA, and high ecDNA copy numbers impose a substantial fitness cost on tumor cells^13^. We propose that suppression of KRAS-stimulated lysosome function by high MYC expression contributes to this fitness cost and thereby drives heterogeneity in MYC expression: cells with high MYC levels experience reduced lysosome function and a growth disadvantage, yet at the same time protect neighboring cells with lower MYC levels from immune eradication. Mathematical modeling confirms that this mechanism selects for tumors with highly variable MYC levels and indicates that heterogeneity in MYC expression provides a net growth advantage to the entire tumor.

Finally, we note that our data have direct therapeutic implications. Because MYC associates with MIZ1 independently of MAX^48^, targeting MYC/MIZ1 complexes and the downstream attenuation of lysosome function-could selectively disrupt the immune evasive MYC activity while sparing essential MYC/MAX functions. MIZ1 contains a structured POZ/BTB domain that can serve as a docking site for small-molecule ligands, and ligands have been identified for MIZ1 and its heterodimeric partner BCL6. Additionally, degraders using BCL6 ligands are in clinical exploration^49,50^. Like MYC/MIZ1, BCL6 is a transcriptional repressor that has been successfully reprogrammed by bivalent molecules to activate transcription^51,52^. Conceptually similar molecules that relieve MYC/MIZ1-mediated transcriptional repression could reverse MYC-dependent immune suppression and offer therapeutic potential for patients with pancreatic cancer.

## Resource availability Lead contact

Further information and requests for resources and reagents should be directed to and will be fulfilled by the lead contact, Martin Eilers.

## Materials availability

No unique reagents were generated for this study.

## Data and code availability

- All new sequencing datasets have been uploaded to GEO under GSE319383 for the bulk RNA-sequencing and GSE319988 for the scRNA-sequencing. Reviewer tokens will be provided upon request. The SILAC mass spectrometry proteomics data have been deposited to the ProteomeXchange Consortium via the PRIDE partner repository with the dataset identifier PXD080328. Publicly available RNA-Seq and ChIP-Seq datasets downloaded include GSE154903, E-MTAB 8792, and GSE64425.
- The agent-based model used to generate Fig. 7E has been uploaded as a stand-alone, dependency free HTML file here: https://doi.org/10.5281/zenodo.21476272. All simulations are deterministic and parameters and select seeds have been reported in Supplementary Table 5.
- Any additional information required to reanalyze the data reported in this paper is available from the <u>lead contact</u> upon request.

## Authors’ contributions

B.K., T.K., G.M., A.R., and M.E. participated in research design. B.K., G.M., M.J., A.C. performed flow cytometry and scRNA sequencing experiments. T.K. performed all bioinformatic analysis. W.S., J.B., and B.W. performed and analyzed mass spectrometry. B.K., A.G.W. and G.B. applied and conducted orthotopic transplantation. C.P.A. performed the sequencing. B.K., A.A., S.C. and M.V. designed and conducted *ex vivo* analysis of T cells. S.B. designed knockout of *B2m*. B.K., L.N. and C.S.-V. performed high-content immunofluorescence experiments. S.M, F.Z., M.F.R generated the MIZ1^fl/fl^ mice, F.Z and D.S. crossed and bred the MIZ1 mouse models. S.H. characterized MIZ-KO cell line. C.S. provided unpublished data. M.F.R. reviewed and edited the paper. B.K., T.K., and M.E. conceptualized the study and wrote the paper.

## Acknowledgments

This work was supported by grants from the European Research Council (SENATR ERC #101096948 to M.E.), the German Cancer Aid (#70114538 to M.E.), the Mildred Scheel Junior Research Center Program (to B.K., A.R., G.B. and M.V.), the German Research Foundation (EI 222/24-1 to M.E.), the KOODAC team supported by the Cancer Grand Challenges partnership funded by Cancer Research UK, the French National Cancer Institute (INCa) and the Children Cancer Free Foundation (Kika), the National Institute of Health CA096832 and CA21765 (MFR) and the American Lebanese Syrian Associated Charities (ALSAC). Operetta microscope was funded by the German Research Foundation (INST 93/1023-1-FUGG). Cytek Auroa 3L was funded by the German Research Foundation 549250077 (INST 93/1162-1-FUGG). Incucyte was Funded by the Deutsche Forschungsgemeinschaft (DFG, German Research Foundation) 506524377 (INST 93/1112-1-FUGG). The authors thank Sarah Hess, Tobias Roth, Barbara Bauer, Ciler Maden, and Marion Krafft for technical support. We thank Rachel Levine from the Center for Advanced Genome Engineering (St. Jude Children’s Research Hospital) for help in the knock-out and Sarah Robinson for genotyping and colony management.

## Declaration of generative AI- and AI-assisted technologies in the manuscript preparation process

During the preparation of this work, the author(s) used PerplexityPro for deep research. Claude code was used to help create the dashboard version of the mathematical modeling, and the code was manually reviewed. The author(s) reviewed and edited the output as needed and take full responsibility for the content of the published article.

## Supplementary Figure Legends

**Supplementary Figure 1:**
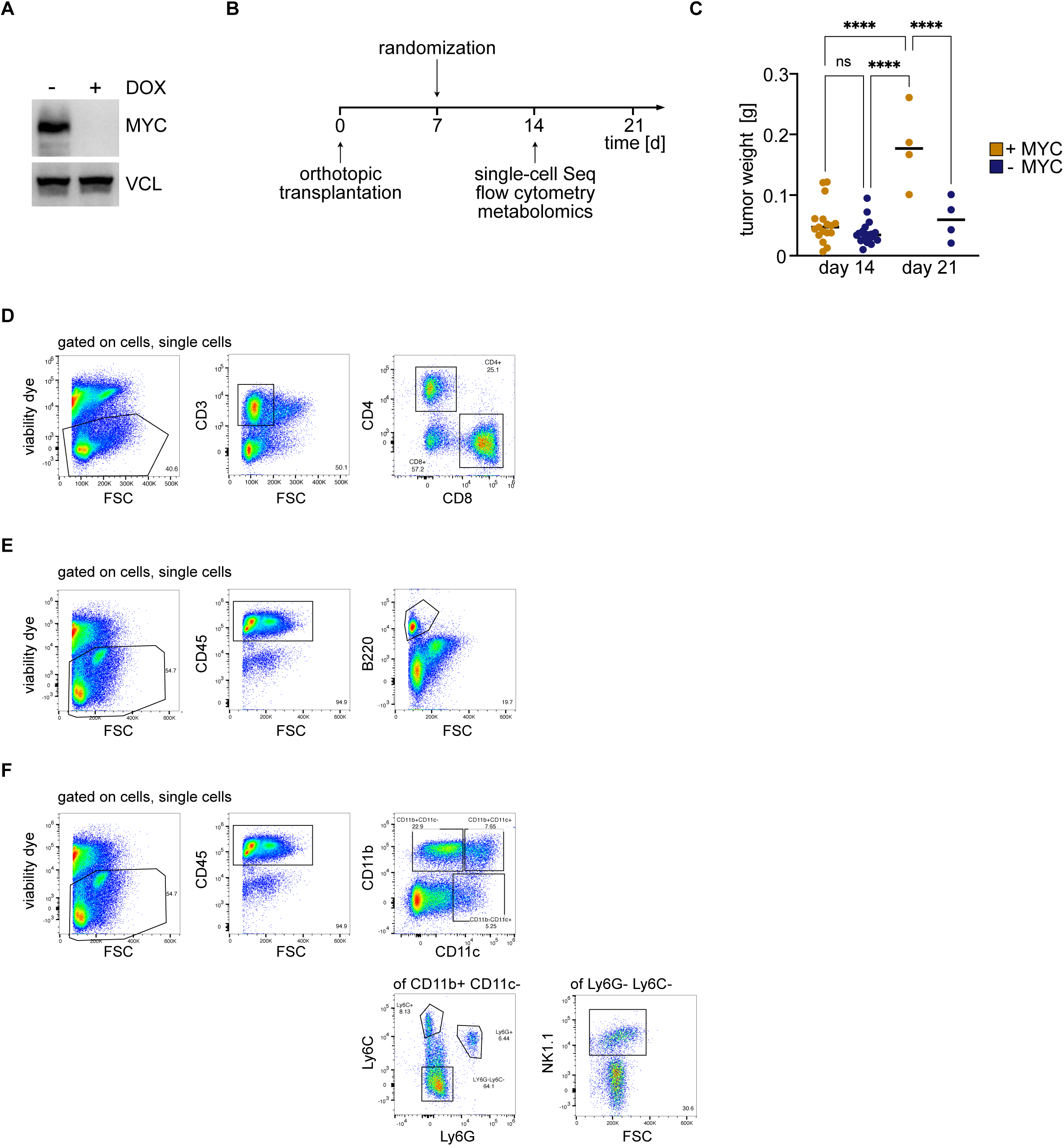
(A) Timing of transplant experiments and analyses. Mice were randomized on day 7 and treated with doxycycline-containing or control food from day seven onwards. Samples for flow cytometry, mass spectrometry and single cell sequencing were collected at day 14. (B) Immunoblot of KPC cells with doxycycline-inducible, shRNA mediated depletion of MYC (1 µg/mL, 48 h). Vinculin was loaded as control. (C) Weight of tumors at day 14 and day 21 (N = 17, 18, 4; error bars: SD; ordinary one-way ANOVA). (D) Gating strategy for flow cytometry of T cells. (E) Gating strategy for flow cytometry of B cells. (F) Gating strategy for flow cytometry of myeloid cells.

**Supplementary Figure 2:**
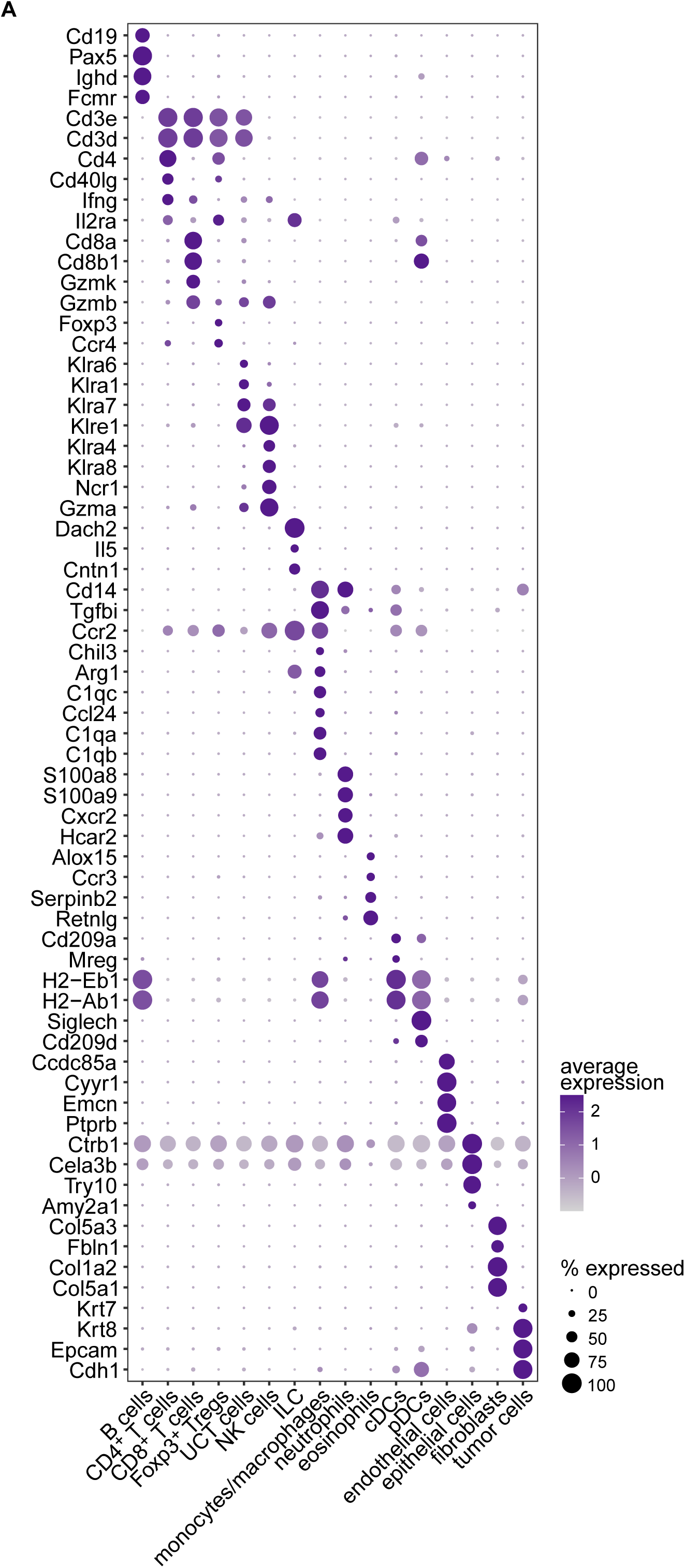
(A) Dot plot of cluster markers corresponding to the different annotated compartments from the single cell RNA-Sequencing of + MYC and - MYC tumors.

**Supplementary Figure 3:**
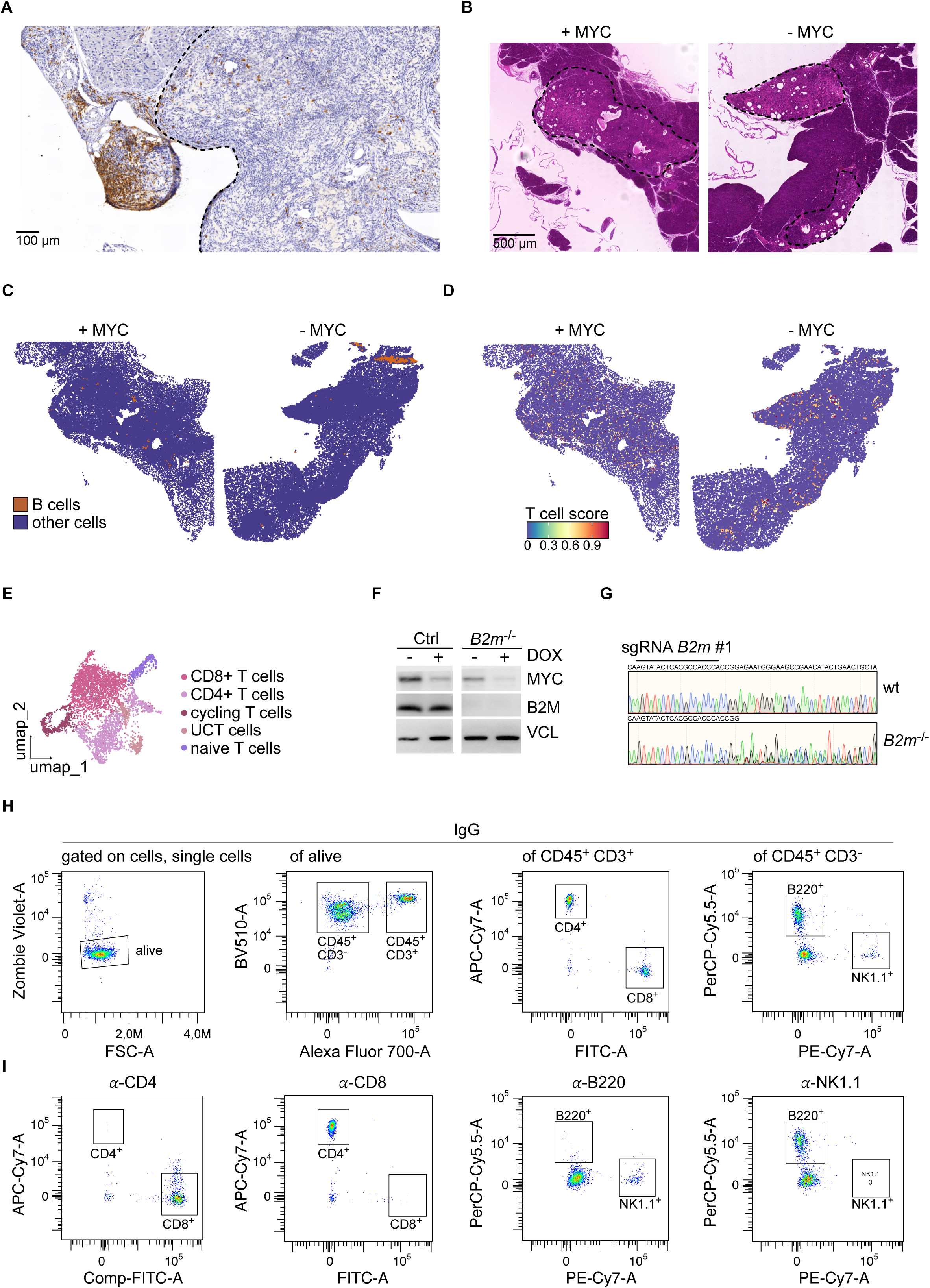
(A) DAB-staining of selected tissue area after depletion of MYC for 7 days stained using B220-antibody; the dashed line indicates tumor circumference. (B) H&E staining of + MYC and - MYC tumor sections. Tumor area highlighted with dashed line. Treatment with doxycycline to deplete MYC was for 48 h. (C) Spatial transcriptomic feature plot highlighting B cells predominantly in the tumor periphery. MYC was depleted by treatment with doxycycline for 48 h. Same slide sections as (B) were used for spatial transcriptomics (N = 1 per condition). (D) Visualization of T cell signature in the same KPC-cell derived tumors. (E) UMAP of T cells derived from 5’ scRNA sequencing. (F) Immunoblot of KPC cells expressing doxycycline-inducible shRNAs targeting MYC. *B2m* was deleted using CRISPR/Cas9. Vinculin serves as loading control. (G) Sanger sequencing of locus of the *B2m* gene that was targeted using CRISPR/Cas9. Sequence downstream of the sgRNA-binding region shows heterozygous mutations, leading to deletion of B2m expression. (H) Gating strategy for flow cytometry of PBMCs of mice with depletion of specific lymphocytes. Example shows PBMCs from mice treated with IgG. (I) Flow cytometry of PBMCs of tumors treated with either *α*-CD4, *α*-CD8, *α*-B220, or *α*-NK1.1 antibodies for selective lymphocyte depletion.

**Supplementary Figure 4:**
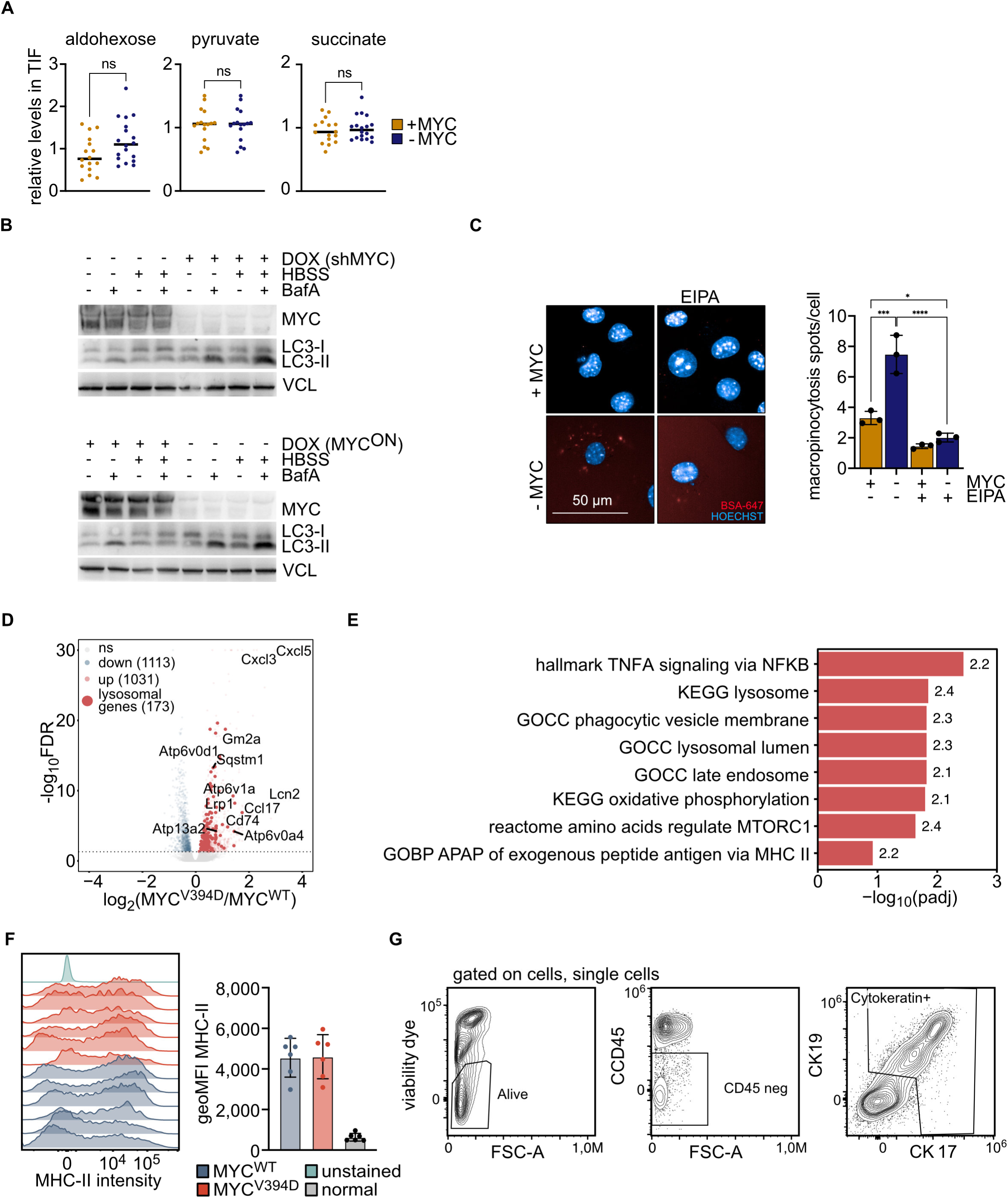
(A) LC-MS analysis of interstitial fluid from tumors before and after MYC depletion (N = 17, 18; unpaired, two-sided t-test). (B) Immunoblots of MYC and LC3 in KPC cells grown in normal conditions (HPLM) or under starvation (HBSS), treated with BafA as indicated. Top panel shows doxycycline induced depletion of endogenous MYC. Bottom panel shows doxycycline induced depletion of endogenous MYC and exogenous expression of MYC^WT^. Vinculin is loading control. (C) Immunofluorescence of KPC cells (+/- MYC) treated with BSA-Alexa647 for 1 h. Where indicated, cells were treated with 25 μM EIPA (5-(N-ethyl-N-isopropyl)amiloride; inhibitor of macropinocytosis) for 24 h. Scale bar indicates 50 µM. Bar plot shows number of Alexa647-spots (N = 3; error bars: SD). (D) Volcano plot showing differentially expressed genes in mRNA-sequencing of KPC cells expressing MYC^V394D^ vs MYC^WT^. Genes that pass FDR < 5% are statistically significant. Lysosome related upregulated genes have been highlighted. (E) Bar plot showing upregulated pathways following a gene set enrichment analysis (GSEA) of the RNA-sequencing in (D). The numbers on the ends of the bars indicate the normalized enrichment score (NES). (F) MHC-II intensity and quantification in tumor cells expressing either MYC^WT^ or MYC^V394D^ compared to normal cells *in vivo* (N = 6). (F) Gating strategy of MHC II presentation on tumor cells.

**Supplementary Figure 5:**
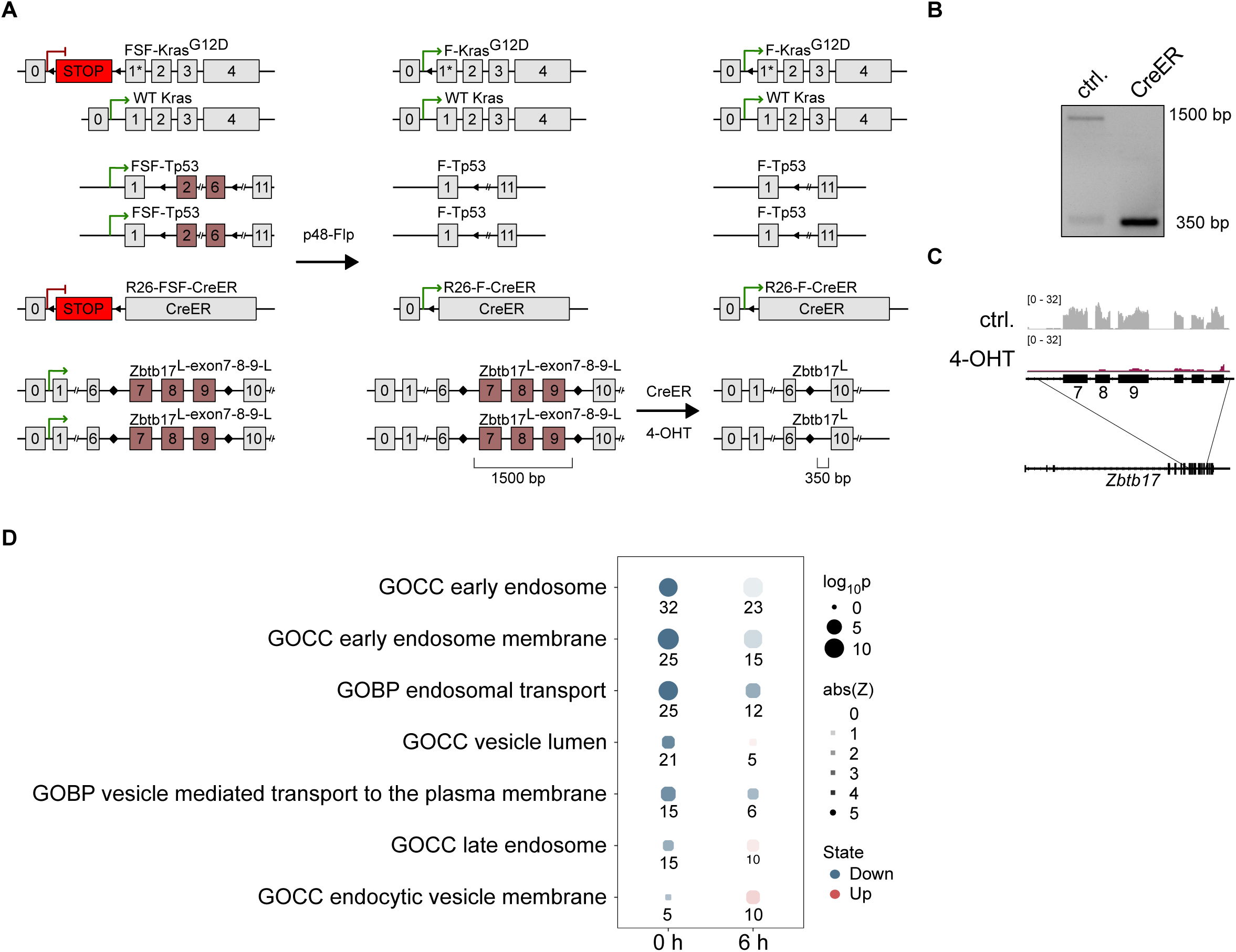
(A) Scheme of KPC mouse model with a conditional allele of MIZ1 (*Zbtb17*). (B) PCR showing deletion of exon 7, 8 and 9 of the MIZ1 gene after induction of the Cre-recombinase. (C) Browser track of RNA sequencing indicating deletion of exons 7, 8 and 9 and strong reduction of expression of downstream exons. (D) Over-representation analysis of proteins differentially labeled in -MIZ1 vs +MIZ1 cells. Size of dots correlate with significance; intensity corresponds to Z-score that indicates direction and strength of enrichment in said direction. Number indicates number of proteins in the pathway differentially labeled.

**Supplementary Figure 6:**
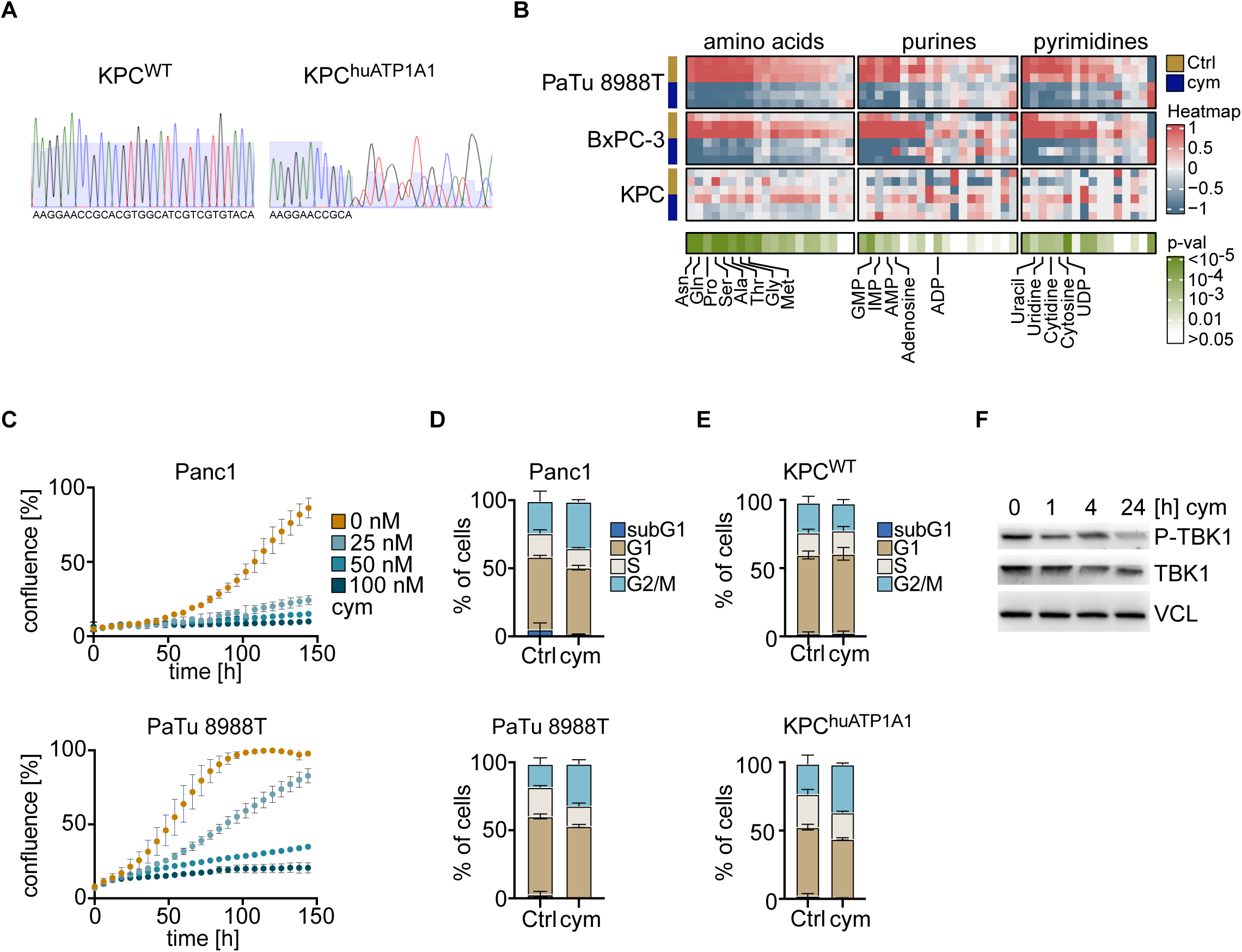
(A) Sanger sequencing of KPC cells with CRISPR/Cas9 mediated targeting of the *Atp1a1* locus shows an insertion and frame shift downstream of the sgRNA target site. (B) LC-MS analysis of intracellular metabolites in human and murine PDAC cell lines treated with 100 nM cymarin for 24 h (N = 3). (C) Growth curve of human PDAC cell lines treated with the indicated concentrations of cymarin (N = 3; error bars: SD). (D) Cell cycle analysis of human PDAC cell lines using propidium iodide staining. Cells were treated for 24 h with 100 nM of cymarin (N = 3; error bars: SD). (E) Cell cycle analysis KPC^WT^ and KPC^huATP1A1^cells using propidium iodide staining. Cells were treated for 24 h with 100 nM of cymarin (N = 3; error bars: SD). (F) Immunoblot of KPC^huATP1A1^ cells treated with 100 nM cymarin for 4 h and 24 h. Vinculin serves as loading control.

**Supplementary Figure 7:**
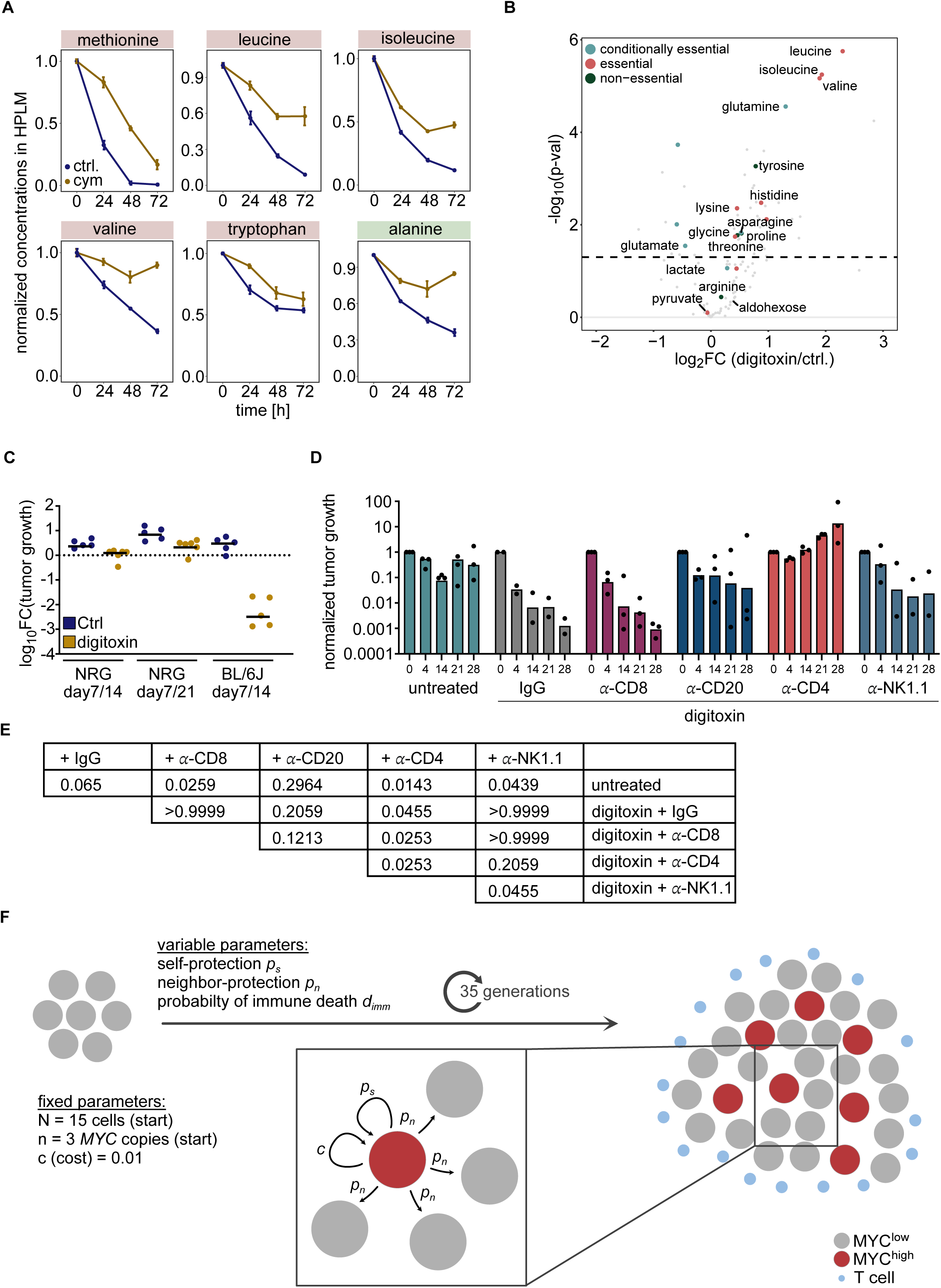
(A) Time-course analysis of LC-MS of supernatant of KPC^huATP1A1^ tumor cells treated with 100 nM cymarin, normalized to respective metabolite levels in the medium (N = 3; error bars: SD). (B) Volcano-plot of LC-MS analysis of tumor interstitial fluid comparing unperturbed KPC^huATP1A1^ tumors and KPC^huATP1A1^ tumors treated for 12 h with digitoxin (N = 9). (C) Quantification of luciferase signal after treatment of tumor bearing C57BL/6J mice and NRG mice with digitoxin for 7 and 14 days. Luciferase signal is normalized to day 7. (D) Quantification of luciferase signal after treatment of tumor bearing C57BL/6J mice with digitoxin and IgG (N = 2), *α*-CD8 (N = 3), *α*-CD20 (N = 3), *α*-CD4 (N = 3) and *α*-NK1.1 (N = 2) antibody. Luciferase signal is normalized to day 5 after transplantation. Mice were treated with digitoxin after 7 days of engraftment. (E) P values of comparisons of the overall survival between different treatments in Figure 7D (Gehan-Breslow-Wilcoxon test). (F) Scheme for mathematical modeling predicting effect of bystander protection by *MYC* high tumor cells.

**Supplementary Data 1**: Html dashboard for the agentic AI-aided modeling, simulating bystander protection conferred by MYC high cells. Input hyperparameters include the coefficients for self-protection, neighbor protection, killing probability, and random seed. Fixed parameters are number of cells in initial lattice = 15, cost per copy of ecDNA = 0.01, number of generations of replications simulated = 35. Toggle on the right lets user choose between an independent model for protection and an additive model. Three panels are generated, first assuming no immune response, second with only a selfish protection of MYC high cells and the third where MYC high cells confer protection on both themselves and their neighbors. Barplots below panels indicate ecDNA distribution. Simulation results can be exported as PDFs. The dashboard can be downloaded using https://doi.org/10.5281/zenodo.21476272 and accessed on any web browser.

## Methods

### Ethics, study approval and animal maintenance

All orthotopic transplantations were approved by the government of Unterfranken under the protocol numbers RUF 55.2–2532-2-2103, RUF 55.2–2532-2-1026, RUF 55.2–2532-2-148, and RUF 55.2–2532-2-1419. Mice were closely monitored by authors, facility technicians and an external veterinarian. MIZ1^fl/fl^ mice were housed in an accredited facility of the Association for Assessment and Accreditation of Laboratory Animal Care International (AAALAC) in accordance with National Institute of Health guidelines. The Animal Care and Use Committee (ACUC) of St Jude Children’s Research Hospital approved all procedures concerning the generation of MIZ1^fl/fl^ mice. MIZ1^fl/fl^ mice were crossed with KPC mice under the protocol number ROB-55.2-2532.Vet_02-22-149.

### Orthotopic transplantation

Orthotopic transplantation was conducted in male C57BL/6J as previously described ^53^. NRG mice (SOPF NOD-Rag2 tm1-Il2rg tm1/Rj) were received from Janvier. Surgery was performed in a biological safety cabinet, and mice were kept in IVC cage. For antibody-mediated depletion tumor size was determined 5 days after transplantation. Mice were randomized according to IVIS signal. For depletion of lymphocytes, antibodies were injected in 200 µL sterile PBS 5 days, 7 days and 21 days after transplantation with the following amounts: 100 µg α-CD8b (Lyt 3.2, BioXCell), 300 µg α-CD4 (GK1.1, BioLegend), 120 µg α-NK1.1 (PK136, BioLegend), 250 µg α-CD20 (SA271G2, BioLegend), and 300 µg IgG (BioLegend). For the depletion of macrophages antibodies were daily injected from day 5 to day 28 in 200 µL PBS with the following amounts: 300 µg α-CD115 (AFS98, BioLegend) and 300 µg IgG (BioLegend). Depletion of immune cell populations was confirmed with flow cytometry. Mice received doxycycline or digitoxin from day 7 onwards.

### Cell culture

KPC cells ectopically expressing of MYC^WT^ and MYC^V394D^ have been described ^18^. Panc1, PaTu 8988T and murine KPC cell lines were grown in DMEM medium containing 10% fetal calf serum (FCS) and 1% penicillin/streptomycin. BxPC-3 cells were grown in RPMI-1640 supplemented with 10% FCS and 1% penicillin/streptomycin. The identity of human cell lines was verified by STR profiling. All cell lines were routinely verified to be Mycoplasma-free. Cells were kept at 37 °C and 5% CO2. Where indicated, cells first adapted to HPLM medium (Thermo) containing 10% fetal calf serum (FCS) and 1% penicillin/streptomycin (called TPLM). Experiments were then performed in TPLM medium. To mimic increase in available amino acids in the TIF after treatment of mice with digitoxin we added an amino acid cocktail (final concentration (without metabolites from HPLM): 2.25 mM glutamine, 0.8 mM glycine, 0.15 mM histidine, 0.63 mM leucine, 0.66 mM valine). For generation of KPC^huATP1A1^ cells were lentivirally infected with a construct expressing ATP1A1 under the SV40 promotor. Cells were subsequently infected with CRISPR V2 expressing an sgRNA targeting the murine *Atp1a1* (CACCGGTACACGACGATGCCACGTG).

### Interstitial fluid and plasma preparation

To collect plasma, mice were bled from the cheek. Blood was collected in EDTA-containing tubes and centrifuged at 4 °C for 10 min at 850 xg. Supernatant was moved to a new tube and centrifuged for 20 min at 3,000 xg. Supernatant was moved to a new tube and flash frozen. For the collection of interstitial fluid, tumor tissue was dried on Whatman paper from PBS and placed on 20 µM cell strainers and 10 µL PBS were added on top. Tissue was centrifuged for 10 min at 4 °C and 150 xg. Flow through was transferred to a new tube and flash frozen. Interstitial fluid was diluted using water to adjust volume according to the weight of the tumors.

### Generation of Miz1 (Zbtb17) conditional knock-out mice

The knock-out was performed using two chemically modified single guide RNAs (sgRNAs, Synthego) [SNP298.g5 and SNP299.g12] and two single-stranded oligodeoxynucleotides (ssODNs) [SNP298.sense and SNP299.sense] to introduce a LoxP site 5’of coding exon 5 of mZbtb17 (Miz1) and a LoxP site 3’ of coding exon 7. Upon Cre expression, exons 5-7 were recombined resulting in the deletion of Miz1/Zbt17. The two gRNAs (10 ng/ul each), the two ssODNs (10 ng/ul each) and Cas9 protein (40 ng/ul) were injected into C57BL/6NJ single cell embryos. Three founder mice, carrying the two LoxP sites on the same allele were obtained and used to establish three independent Miz1-floxed mouse lines #10, #17 and #18. Experiments were performed using line #10.

sgRNA spacers: SNP298.g5 (cccugccugcgcacgacaca), SNP299.g12 (uccuaggauggucccccgga), ssODNs (both contain a BamH1 site and a LoxP site: SNP298.sense(agtgtggaagcatcctgacacagtcctggagcttcctgctcagccctgcctgcgcacgacATAACTTCGTATAATGTATGCTATACGAAGTTATGGATCCacatggtgtgccatgttgcccagaaacctcacaaggacttggttggctgaaaatgcccac) SNP299.sense(tggagctcccgcgatagcacacaacagccagggacagggactctcctaggatggtcccccGGATCCATAACTTCGTATAATGTATGCTATACGAAGTTATggagggctgtggttgcttttccttccagtgctgggacctgggcttctatcaggaatagag)

Mice were crossed with KPC mouse model (Ptf1a-Flp, FSF-Kras^G12D^, Tp53^fl/fl^, FSF-R26-CreER). We used these primers to validate deletion of the exons: TCTTCATTGCCAGTCCCCAT, TGGATCTTCAAGTGGGCCTT

### Immunoblotting

Immunoblotting was performed as previously described in ^53^.

### Immunofluorescence

To stain lysosomes, living cells were stained with Hoechst in full medium for 10 min at 37 °C, 5% CO2. Hoechst-containing full medium was removed and replaced by full medium containing 50 nM Deep Red Lysotracker. Cells were incubated for 30 min at 37 °C and 5% CO2. The 96-well plate was analyzed immediately afterwards using an Operetta System high Content Imaging System and analyzed using the Harmony software.

Tumors were sectioned at 10 μm thickness in the Cryostat machine and transferred to glass slides. Sections were then frozen at -80°C until IF staining. 10 μm frozen sections were then fixed in ice-cold acetone:methanol 1:2 mix for 2 minutes, blocked with 5% goat serum 2% bovine serum albumin (BSA) in phosphate buffered saline (PBS), and incubated with primary anti-mouse antibodies at 4 °C overnight. Alexa Fluor secondary antibodies were used at a dilution of 1:300. Sections were mounted in ProLong Gold and epifluorescence images were taken using DMi8 Inverted Microscope and analyzed using Fiji.

### Macropinocytosis assays

To stain pinocytosis cells were stained with Hoechst in full medium for 10 min at 37 °C, 5% CO2. Hoechst-containing full medium was removed and replaced by HPLM containing 250 µg/mL BSA-Alexa647. Cells were incubated for 1 h at 37 °C and 5% CO2. Staining medium was replaced by HPLM. The 96-well plate was immediately afterwards measured using the Operetta System. The analysis was performed using the Harmony software.

### Isolation and differentiation of macrophages

Femur and tibia of mice were isolated and tissue removed completely. Femur ends were cut and bone marrow flushed with HPLM full medium with a 21-gauge needle. Bone marrow was passed through a 70 µM cell strainer. Cells were cultivated in full HPLM medium with 10 ng/mL M-CSF. Medium was replaced every second day. Sample preparation and flow cytometry analysis is described below.

### Purification and differentiation of T and NK cells from splenocytes

Mice were euthanized by CO₂ inhalation or cervical dislocation. Single-cell suspensions from lymphoid organs (spleen and lymph nodes) were generated by mechanical dissociation. Briefly, tissues were placed onto a 70-µm cell strainer fitted onto a 50-ml falcon tube and gently disrupted using the plunger of a 1-ml syringe. For lymph nodes, which do not contain erythrocytes, the strainer was pre-wetted with PBS and subsequently rinsed multiple times with PBS to maximize cell recovery. For splenic samples, red blood cells were removed by incubation with two sequential additions of 5 ml red blood cell (RBC) lysis buffer for 1–3 min. Lysis was terminated by dilution with PBS. Cell suspensions were kept on ice and centrifuged at 1200 rpm for 5 min at 4 °C. Pellets were resuspended in PBS, and viable cells were counted prior to further use. Murine CD8^+^ T cells were isolated from pooled spleen and lymph node cell suspensions of male and female mice using a negative selection approach. Purification was performed with the MojoSort™ Mouse CD8 T Cell Isolation Kit or CD8a^+^ T cell Isolation Kit, mouse for CD8^+^ T cells, and with the CD4^+^ T cell Isolation Kit, mouse for CD4^+^ T cells, according to the manufacturer’s instructions. Isolated cells were directly used for in vitro culture experiments. Cells were cultured under sterile conditions at 37°C in a humidified incubator with 5% CO₂. HPLM media was used to mimic the tumor microenvironment.

For T cell activation, delta-surface culture plates were pre-coated with 12 µg/mL polyclonal anti-hamster IgG for at least 2 h at 37 °C and subsequently washed once with PBS. CD8^+^ T cells were seeded at 1.6 × 10⁶ cells in 2 mL HPLM medium in 12-well plates. Cells were activated for 48 h with plate-bound anti-CD3 (0.5 µg/mL) and soluble anti-CD28 (1 µg/mL). Following activation, CD4^+^ and CD8^+^ T cells were differentiated under subset-specific conditions for an additional two and four days, respectively. Effector-like CD8^+^ T cells (Teff) were cultured in the presence of 10 ng/mL recombinant human IL-2. Memory-like CD8^+^ T cells (Tmem) were generated using 10 ng/mL IL-7 and 10 ng/mL IL-15. Naïve (Th0) cells were cultured in the presence of 2.5 µg/ml anti-IFN-g, 2.5 µg/ml anti-IL-4, and 10 ng/mL IL-2. Th1 cells were cultured with 2.5 µg/ml anti-IL-4, 10 ng/ml IL-12, and 10 ng/ml IL-2. Th2 cells were differentiated in the presence of 2.5 µg/ml anti-IFN-g, 50 ng/ml IL-4, and 10 ng/mL IL-2. Induced Treg (iTreg) cells were generated with 2.5 µg/mL anti-INF-g, 2.5 µg/ml anti-IL-4, and 5 ng/mL TGF-b1. Conventional Th17 (Th17c) cells were cultured in the presence of 2.5 µg/ml anti-IFN-g, 2.5 µg/ml anti-IL-4, 20 ng/mL IL-6, and 5 ng/mL TGFb1, whereas pathogenic Th17 (Th17p) were cultured with 2.5 µg/mL anti-IFN-g, 2.5 µg/mL anti-IL-4, 20 ng/mL IL-6, 20 ng/mL IL-1b and 20 ng/mL IL-23. To ensure optimal growth and differentiation, cultures were split daily from day 2 onward and maintained at a density of 0.5 × 10⁶ cells/mL. NK cells were extracted from splenocytes using NK cell isolation kit. NK cells were cultivated in HPLM and stimulated with IL15 (150 ng/mL) and IL2 (100 ng/mL) for 6 days. NK cells were treated with amino acids as indicated above. Medium was refreshed every second day. T and NK cell culture was supplemented with 55 nM 2-Mercaptoethanol.

### Flow cytometry

Flow cytometry was performed on cells isolated from tumors as previously described. Briefly, primary tumors were collected from mice, measured using a caliper and weighed to determine the volume and size. Tumors were placed in 500 µL of digestion mixture containing 1 mg/mL collagenase A and D, 2 mg/mL of trypsin inhibitor and 0.4 mg/mL DNase I in digestion media (RPMI supplemented with 2% FBS and 2 mM HEPES). Tumors were then minced into small pieces using a scalpel and incubated at 37°C for 60 min with 1000 rpm agitation in a thermo-mixer. Ethylenediaminetetraacetic acid (EDTA) at a final concentration of 10 mM was then added and cells were passed through a 70 µm strainer prior to immunostaining.

For flow cytometry staining, 1 x 10^6^ cells were stained with fixable viability dye Live/Dead eFluor 780 washed with FACS buffer (0.5% BSA, 2 mM EDTA in PBS) and incubated with Fc receptor blocker for 10 minutes at 4°C, followed by cell surface staining using combinations of fluorescently conjugated antibodies. All antibodies were used at a dilution of 1:100. The cells were then washed in FACS buffer and resuspended in FACS buffer before analysis on an Attune NxT flow cytometer. Peripheral blood was obtained by cheek puncture and collected in microtubes containing EDTA as anticoagulant (50 µL per sample) for flow cytometry analysis of peripheral blood mononuclear cells (PBMCs). Whole blood was incubated for 5 min with red blood cell lysis buffer, followed by washing with phosphate-buffered saline (PBS) and resuspension in PBS. For surface staining, cells were transferred to 96-well plates and incubated for 20 min at 4 °C in 100 µL antibody master mix per well prepared in cold PBS containing viability dye (Zombie Violet, 1:1000), Fc-block (anti-CD16/CD32, 1 µL per sample), and fluorophore-conjugated antibodies at 1:100 dilutions. Following incubation, cells were washed with 100 µL FACS buffer (2 min, 1600 rpm), resuspended in FACS buffer, and acquired on a Cytek Aurora 3-laser spectral flow cytometer. Offline analysis was performed using FlowJo software.

### Flow cytometry-based apoptosis and cell cycle analysis

Annexin V/PI-FACS to determine apoptosis and PI-FACS to determine cell cycle phases were performed as described in ^54^.

### Liquid chromatography-mass spectrometry

For labeling studies KPC cells were cultivated in HPLM medium. We either added 2.5 mM D-glucose-13C6 to the existing 5 mM of glucose or replaced the existing glutamine by 0.55 mM L-glutamine-13C5. We removed HPLM medium, washed cells with prewarmed PBS and replaced this by medium with isotype-labelled metabolites.

We harvested cell pellets washing cells twice with ammonium acetate (154 mM). Cells were scratched, sample adjusted to the same cell number and cell pellet was flash frozen.

Water-soluble metabolites from 3 µL fluid sample (interstitial fluid, plasma or culture supernatant) or cell pellet (1 x 10^6^ cells) were extracted with 500 µL ice-cold MeOH/H2O (80/20, v/v) containing 0.01 μM lamivudine and 1 µM each of D2-glucose, D4-succinate and D5-glycine (Sigma-Aldrich, St. Louis, USA). After centrifugation, the resulting supernatants were evaporated in a rotary evaporator (Savant, Thermo Fisher Scientific, Waltham, USA). Dry sample extracts were redissolved in 100 μL 5 mM NH4OAc in CH3CN / H2O (50/50, v/v). 20 µL supernatant was transferred to LC-vials. Metabolites were analyzed by LC-MS using the following settings: For LC-MS analysis 3 μL of each sample was applied to a SeQuant ZIC-cHILIC (3 μm particles, 100 × 2.1 mm) (Merck, Darmstadt, Germany). Metabolites were separated with Solvent A, consisting of 5 mM NH4OAc in CH3CN/H2O (40/60, v/v) and solvent B consisting of 5 mM NH4OAc in CH3CN/H2O (95/5, v/v) at a flow rate of 200 µL/min at 45 °C by LC using a DIONEX Ultimate 3000 UHPLC system (Thermo Fisher Scientific, Bremen, Germany). A linear gradient starting after 2 min with 100 % solvent B decreasing to 10% solvent B within 23 min, followed by 16 min 10% solvent B and a linear increase to 100% solvent B in 2 min was applied. Recalibration of the column was achieved by 7 min prerun with 100% solvent B before each injection. All MS-analyses were performed on a high-resolution Q Exactive mass spectrometer equipped with a HESI probe (Thermo Fisher Scientific, Bremen, Germany) in alternating positive- and negative full MS mode with a scan range of 69.0 - 1000 m/z at 70K resolution and the following ESI source parameters: sheath gas: 30, auxiliary gas: 1, sweep gas: 0, aux gas heater temperature: 120 °C, spray voltage: 3 kV, capillary temperature: 320 °C, S-lens RF level: 50. XIC generation and signal quantitation was performed using TraceFinder™ V 5.1 integrating peaks which corresponded to the calculated monoisotopic metabolite masses (MIM +/- H+ ± 2 mMU).

### SILAC labelling

Cell line (KPC ZBTB17^fl/fl^) were cultured in complete medium containing either light (Lys0, Arg0), “medium” (Lys4, Arg6) or heavy (Lys8, Arg10) SILAC amino acids for 7 days. Cells were treated with 4-OHT (600 nM) or vehicle for 5 days. Cells were switched to light SILAC medium for a chase period of 0, 1, 2, 4, 6, 9, 18 or 24 h. At each time point, cells were harvested by scraping in RIPA buffer and stored at −80 °C until further processing. Total protein was extracted in RIPA buffer and the protein concentration was determined with the bicinchoninic acid (BCA) assay (Thermo Fisher Scientific). Extracts were diluted to a final concentration of 1 µg µL⁻¹ and 10 µL aliquots were used for downstream processing.

Each aliquot was reduced with 5 mM dithiothreitol (DTT) at 60 °C for 30 min, alkylated with 20 mM chloroacetamide at 37°C for 30 min, and precipitated onto carboxylate coated magnetic beads according to the SP3 protocol ^55^. Reduction, alkylation, precipitation, washing and protease addition were performed on an Agilent Bravo liquid handling system ^56^. Proteolytic digestion was carried out with a LysC/Trypsin mixture (Promega) at a 1:50 (wt/wt) ratio for at 37 °C over night. Digested peptides were resuspended in 1 % trifluoroacetic acid (TFA), desalted using homemade C18 solid phase extraction tips ^57^ and vacuum concentrated. Peptide pellets were stored at −70 °C until mass spectrometric analysis. Peptide concentrations in medium and heavy labeled samples were independently estimated by a short premeasurement on a QExactive Plus mass spectrometer operating in data dependent acquisition (DDA) mode, coupled to a Dionex 3000 UHPLC in trap and elute configuration. Control and OHT treated samples were analyzed either directly or pooled 1:1 to determine M/H ratios. For direct analysis, 1:50 of each sample was injected; for pooled samples, 1:20 was used.

Peptides were injected onto a Vanquish Neo UHPLC system equipped with a 5 mm PepMap Neo C18 trap cartridge and a ProtumLink Extend 15 cm × 75 µm, 1.9 µm C18 analytical column maintained at 50 °C. Chromatographic separation employed a binary solvent system (solvent A: 0.1 % formic acid; solvent B: 80 % acetonitrile, 0.1 % formic acid) over a 24.5 min gradient at a flow rate of 400 nl min⁻¹. Eluted peptides were directly infused into an Orbitrap Astral mass spectrometer operated in positive mode. Background ions were suppressed using field asymmetric ion mobility separation (FAIMS Pro Duo) at a compensation voltage of –45 V, a counter flow gas rate of 4.6 L min⁻¹, and an inner electrode temperature of 100 °C. MSⁱ scans were acquired over a mass range of 380–980 m/z with a resolution of 240 k in the Orbitrap analyzer (AGC target 500 %, maximum injection time 5 ms). Each MSⁱ scan was followed by two 2 m/z window data independent acquisition (DIA) scans at 25 % NCE in the Astral analyzer (AGC target 500 %, maximum injection time 3 ms, scan range 150–2000 m/z). MSⁱ scans were scheduled at a 0.6 s cycle time.

Raw DIA data were were processed with DIA NN^58^ v. 2.3.1. An *in silico* library was first generated in a separate DIA-NN run and used as input for the actual search. Searches were performed allowing a maximum of one missed cleavage, enabling the “reanalyse” option, considering all three SILAC labeling options (light, medium and heavy) as “channels”, specifying the light channel as “fixed-mod” and applying channel run normalisation for pulsed SILAC. The resulting parquet report file was used for all downstream analyses.

Entries were filtered to retain only those with Q.Value < 1 %, Lib.Q.Value < 1 %, and Channel.Q.Value < 1 %. Protein groups were assembled using the MaxLFQ values precalculated by DIA NN. For each time point the heavy/light or medium/light ratios were computed, log₂ transformed, and retained only if at least two of the three replicates yielded valid values. A one sample t test was performed to assess deviation from a ratio of 1; the – log₁₀ p values were plotted against the log₂ ratios to visualise significant changes. The same filtering strategy was applied to the pooled triple SILAC data. Statistical significance was evaluated with a linear regression model implemented in limma^59^ without pairing of replicates. Data analysis and plotting were carried out in Python 3.12. And R 5.4.3.

### Real-time measurement of metabolic activity of cells

Cells were seeded in XFe96 FluxPak. Glycolytic rate assay was performed in the presence of 0.5 μM Rotenone and 0.5 μM antimycin A and/or 100 mM 2-deoxy-D-glucose as indicated. Mito stress assay was performed in the presence of 2 µM Oligomycin, 1 µM FCCP (Mesoxalonitril-4-trifluormethoxyphenylhydrazon) and 0.5 µM Rotenon and Antimycin.

### scRNA sequencing

Tumors were digested and stained as described above before cell sorting. Cell sorting was performed using multicolor FACS (BD FACSAria II). Tumor samples were gated on live cells (Zombie-) and then on CD45+, CD3+ cells to obtain T cells; on CD45+, B220+ cells to obtain B cells; for endothelial cells, on CD45-, Pdpn/Pdgfra-, Cd31+; for stromal cells, on CD45-, Pdpn/Pdgfra+, CD31- and to obtain epithelial cells on Cd45-, Epcam/Ecad+. Antibodies are listed under “Flow Cytometry”

Epithelial, endothelial, and stromal cells were processed using 3’ single-cell RNA sequencing with Chromium Next GEM 3’ v3.1 (10X Genomics). While T cells were processed using 5’ single-cell RNA sequencing with the Chromium Next GEM Single Cell 5’ Kit v2 with Chromium Single Cell V(D)J (10X Genomics). Libraries were quantified by automated electrophoresis (BioAnalyzer, Agilent) and their quality was assessed on a Fragment Analyzer (Agilent). Samples were sequenced using an Illumina NextSeq2000.

### mRNA sequencing

mRNA sequencing was performed as previously described in ^53^.

### Bioinformatic Analysis of 3’ and 5’ scRNA-Seq

Cellranger v9.0.0 ^60^ was used for aligning reads and quantifying transcripts against the mouse transcriptome (refdata-gex-GRCm38-2020-A). Low quality cells that expressed fewer than 200 genes, had more than 15% mitochondrial content, and high ambient RNA content as determined by the algorithm decontX ^61^ were discarded. Seurat v5 ^62^ was used for the downstream analysis including normalization, identification of variable features, and unsupervised clustering. Cluster annotations were performed using markers and verified orthogonally using the R package SingleR ^63^ that predicts cell types using supervised learning. The inbuilt immgen dataset was used as a reference for SingleR predictions. Both Wilcoxon Rank Sum test and DESeq2 on pseudobulked counts were used for differential expression analysis between different conditions in each cell type. Differentially expressed genes that passed an FDR threshold of 5% were deemed statistically significant and pathways over-represented by these genes were identified using hypergeometric tests on pathways from databases such as Hallmark, Reactome, KEGG, GO, Biocarta, and Wikipathways obtained from mouse MSigDB ^64^. Signature scores were computed using the “signed average” expression of log-normalized features for each cell. For the 5’Seq, the Cellranger multi pipeline was used for alignment and reach quantification of the gene expression and the VD(J) libraries. Cells with valid gene expression barcodes after quality control and productive contigs in the TCRB chain were considered for downstream analysis. The R package scRepertoire ^65^ was used for visualization of clones and their expansion.

### Bioinformatic Analysis of bulk RNA-Sequencing

STAR (version: 2.7.10a, ^66^) was used to align the reads against the mm10 genome in the quant mode. The aligned and quantified reads were used as input for normalization and differential expression analysis using the R package DESeq2 ^67^. Genes that passed FDR < 5% (unless specified otherwise) and at least a 1.5-fold change in either direction were deemed statistically significant provided their expression trend was preserved across replicate. Further downstream analysis was conducted both using GSEA on pre-ranked lists as well as over-representation analysis (hypergeometric tests) with statistically significant genes. MCP-counter ^68^ was used to compute cytotoxicity scores in T cells ex vivo. R packages ggplot2 and ComplexHeatmap were used for visualization.

### Bioinformatic Analysis of ChIP-Sequencing

Alignment was done with Bowtie2 and the bam files were read normalized. MACS2 ^69^ was used to call peaks over input and the R package ChIPseeker was used to annotate and identify promoter peaks.

### Bioinformatic Analysis of spatial Transcriptomics analysis

Pre-processing was performed using standard SpaceRanger v3.0.0 pipeline. Seurat RTCD algorithm was used to annotate cell types at the 8um resolution with the scRNA-Seq data from this manuscript as a reference.

### Mathematical modeling of MYC-high cell conferred bystander protection

Cells are assumed to occupy a two-dimensional square lattice of 81x81 sites, following Moore 8-neighborhood and closed boundaries. Every cell has 2 copies of chromosomal MYC.

Let e_i_ by the ecDNA copy number in a cell i. Then,

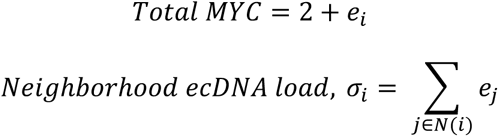

which is the sum over occupied neighborhoods.

The protection arm in this model has two components: a cell can protect itself (p_self_) or be protected by its neighbors (p_pub_).

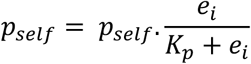

Where a_self_ is the maximum probability of blocking an immune mediated killing. For example, if a_self_ = 0.9, a cell with unlimited ecDNA would block at most of 90% of the killing. K_p_ is the copy number at which the cell gets half the aself as private protection. K_p_ here is set at 8, since PDAC cells typically have 8-15 copies of ecDNA per cell but has been swept to check for consistency of conclusion.

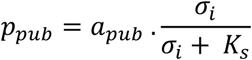

where K_s_ is the total neighborhood ecDNA total at which the protection is half as good as it can be. K_s_ here is set at 20.

Hence, the protection probability is,

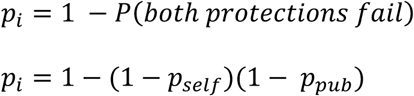

From published work of MYC in PDAC ecDNA^13^, there is a cost/penalty to a cell for carrying ecDNA MYC. If c is the cost of carrying an ecDNA, then cost to a cell i is,

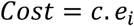

Here, c is arbitrarily fixed at 0.01 but has also been swept to check for consistency.

Finally, if the initial survival probability of a cell is assumed to be 80% and if d_imm_ is the probability of death by immune killing, then survival probability of a cell in each generation is,

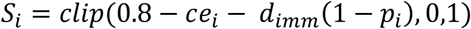

where the clip indicates that survival probability is bound between 0 and 1.

A surviving cell splits into two cells. The chromosomal MYC is faithfully inherited, the ecDNA MYC follows non-Mendelian genetics, so segregates randomly after replicating into 2e_i_ copies.

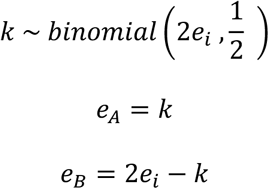

where A and B are the daughter cells of cell *i*.

The model was then programmed using agentic AI and verified with human inspection. Simulations have been initialized with a disc comprising of 15 cells at the lattice center, with each cell initially carrying a random number of copies of MYC ∈ [0,8]. Simulation run for 35 generations, where in each generation, a surviving cell splits into two daughter cells where one daughter cell occupies the position of the parental cell and the other is placed at a uniformly chosed empty adjacent site. If no empty site exists, cell is lost.

In panel 1 of Fig 7E, the immune cells are absent so d_imm_ = 0. In panel 2, the MYC high cells only protect themselves and not their neighbors, so h_pub_ = 0. In panel 3, the MYC high cells protect both themselves and their neighbors so the equation for survival is valid as is. Dashboard of the simulation provided along with manuscript.

### Diagram Generation and Visualization

Diagrams were designed using https://Biorender.com and Affinity Designer 2.

## KEY RESOURCES TABLE

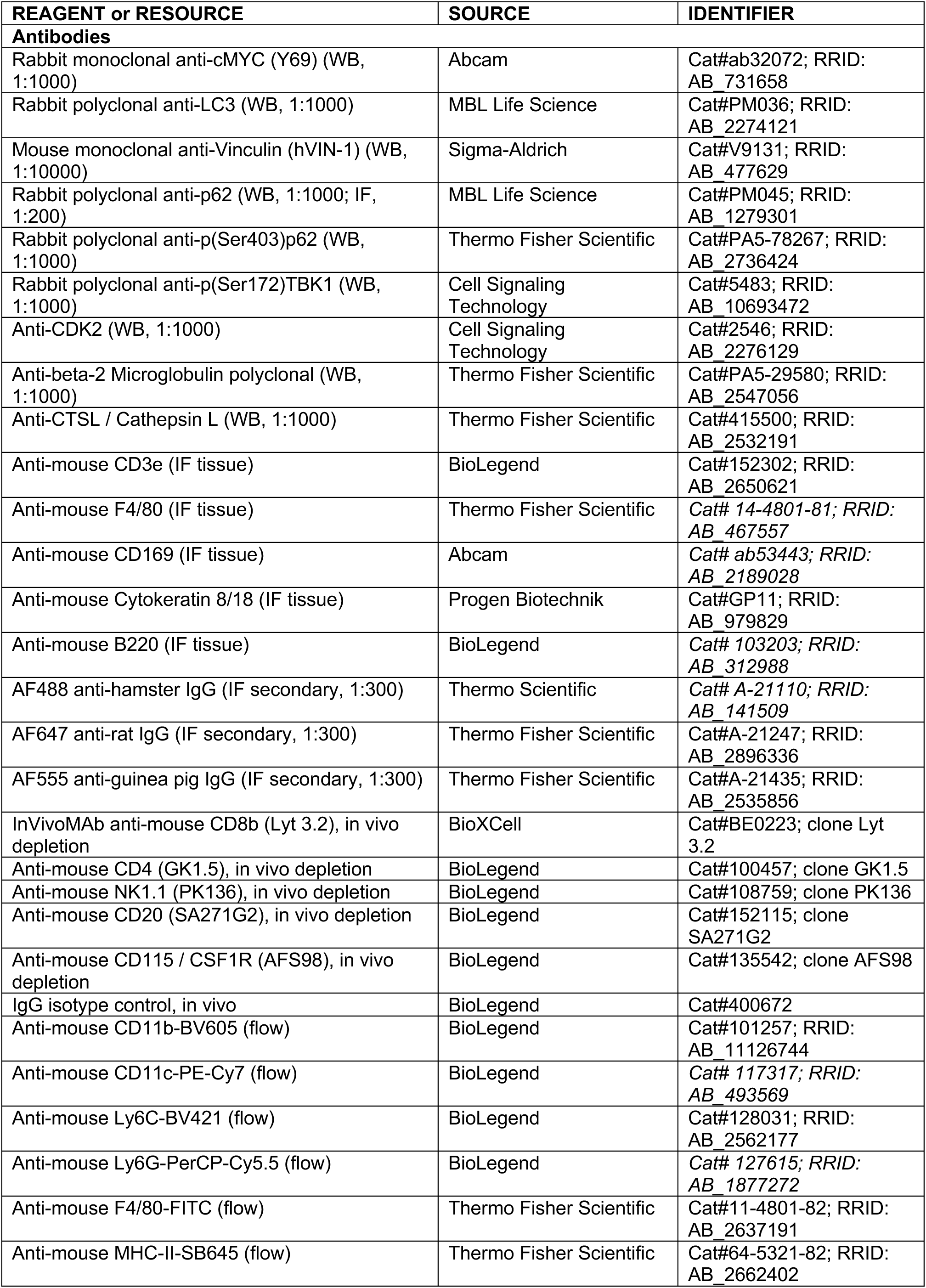
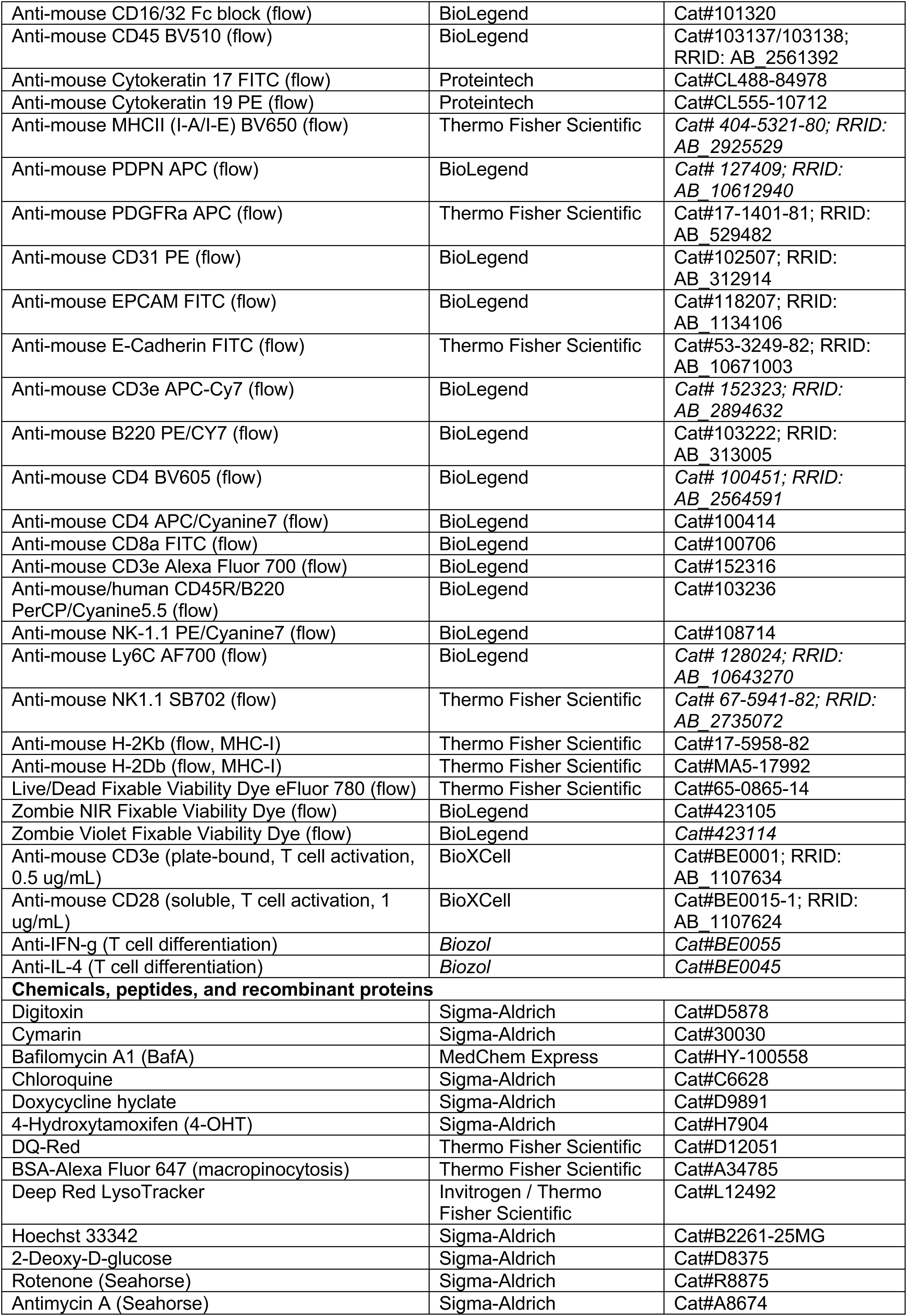
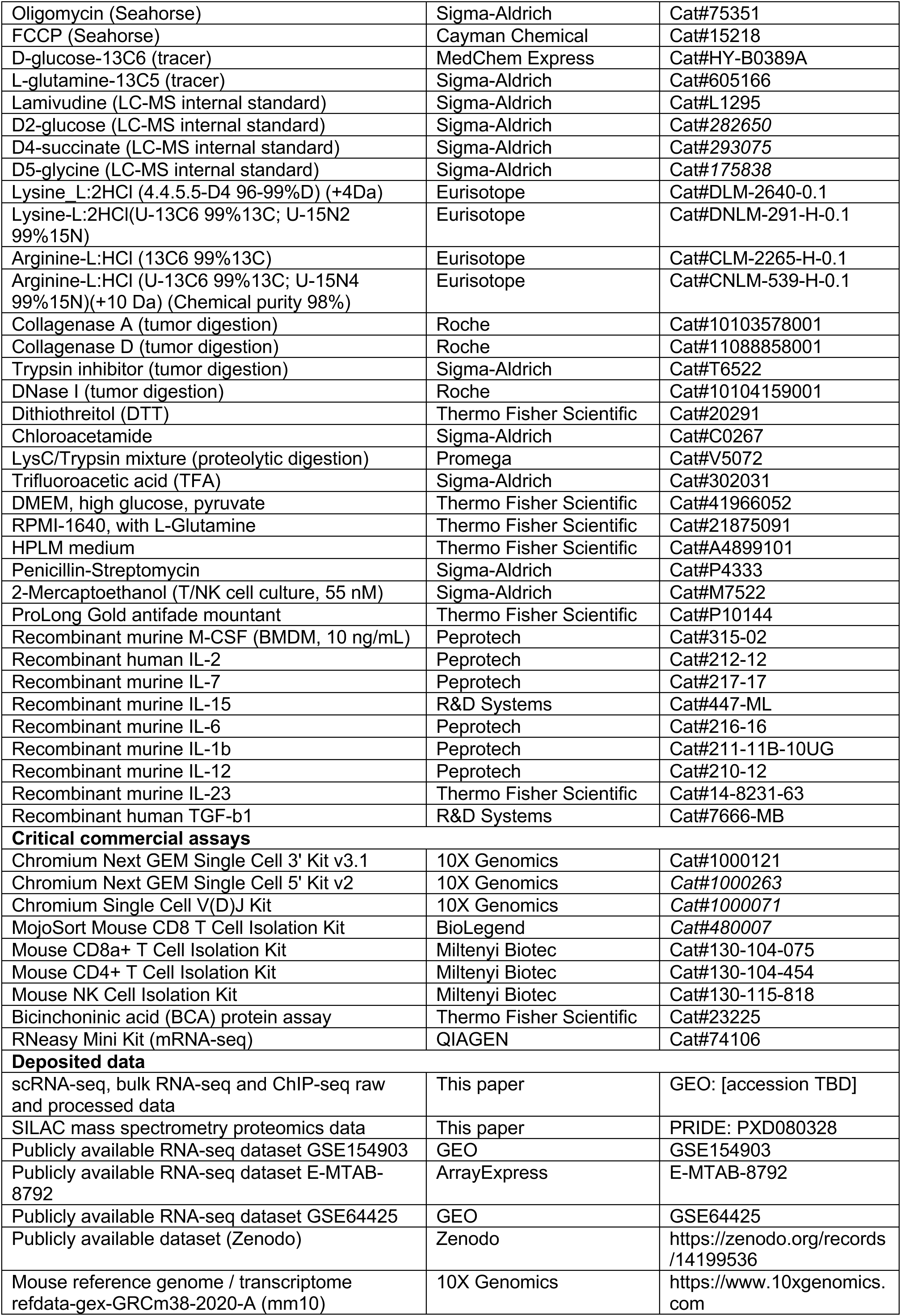
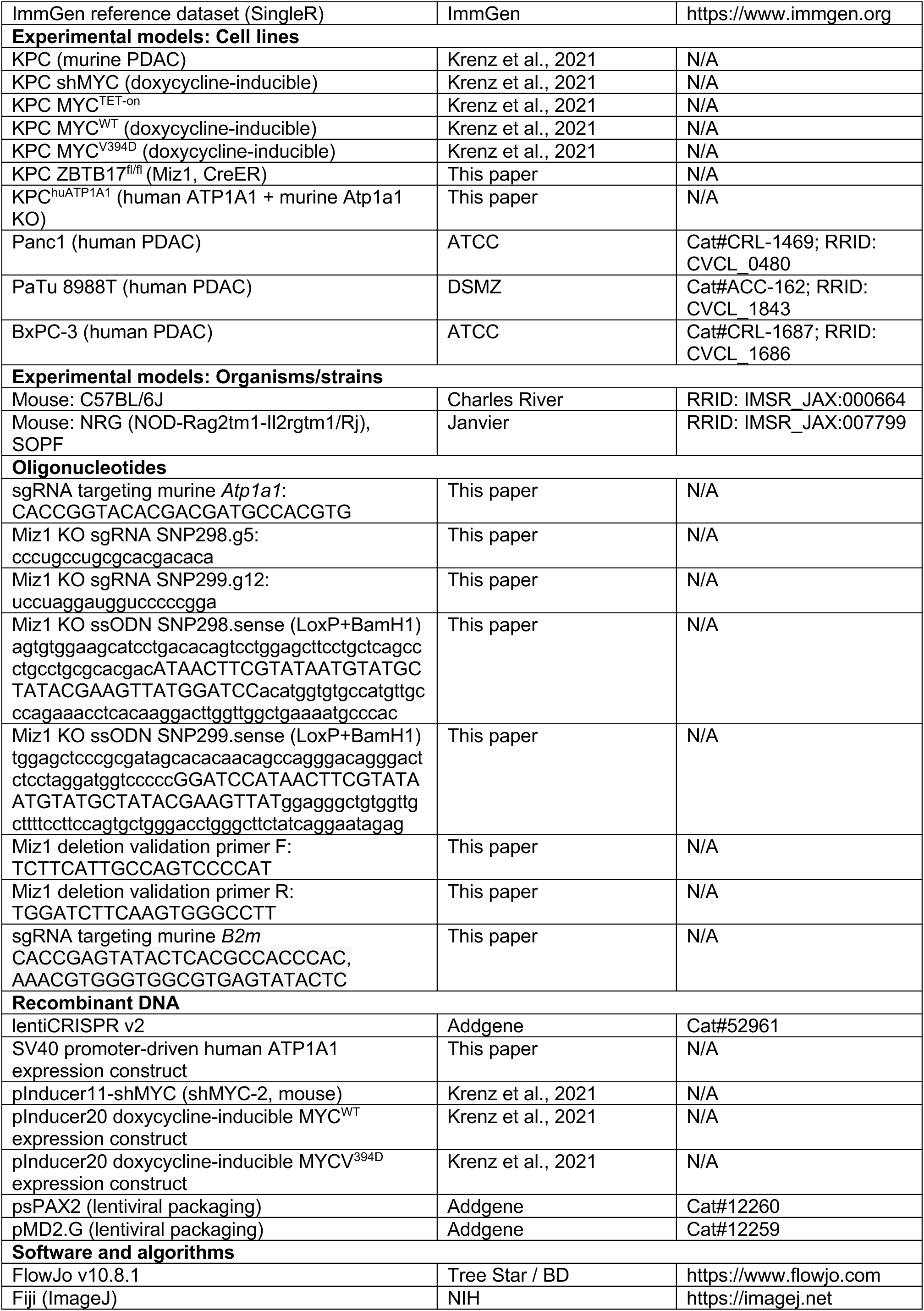
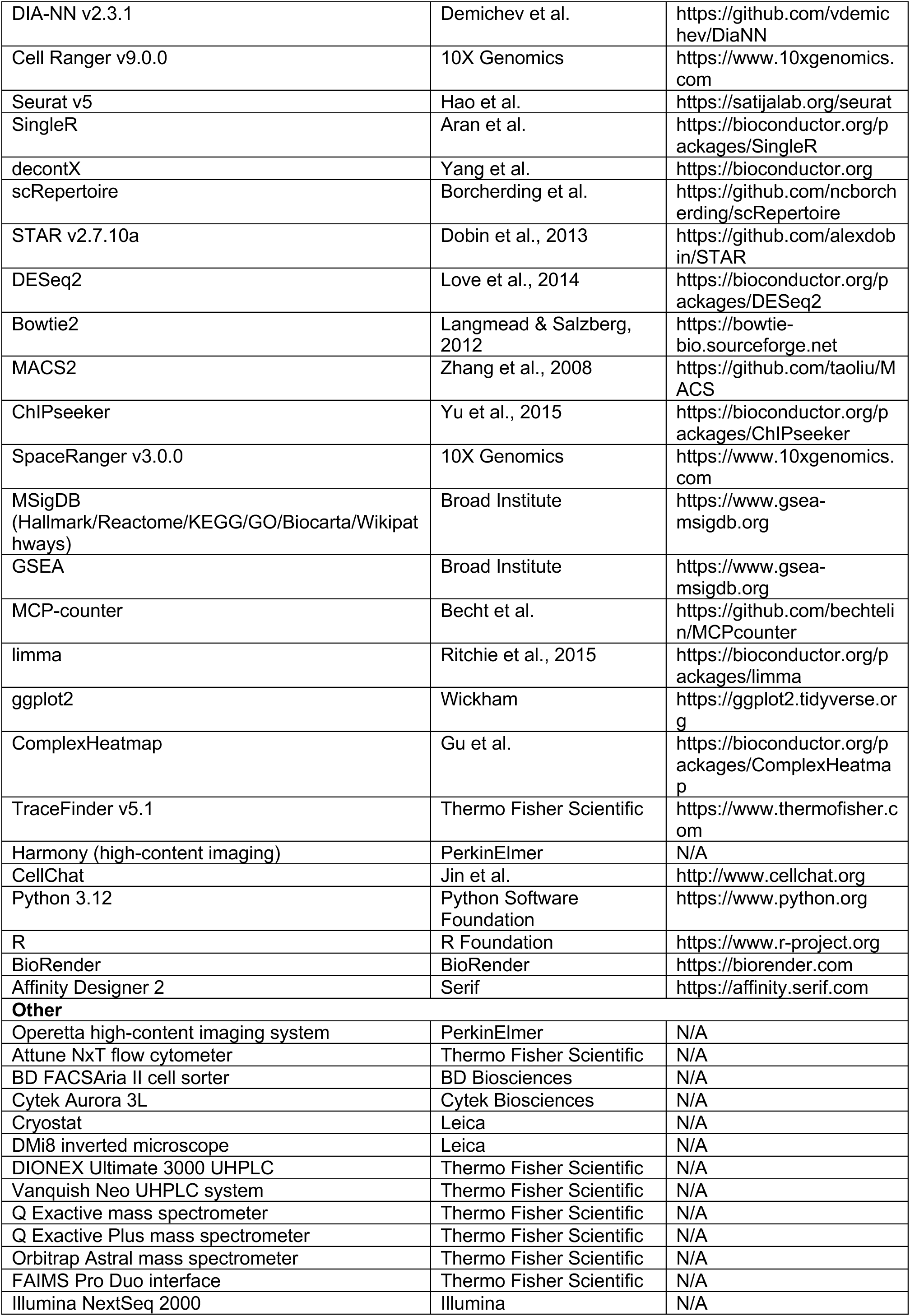
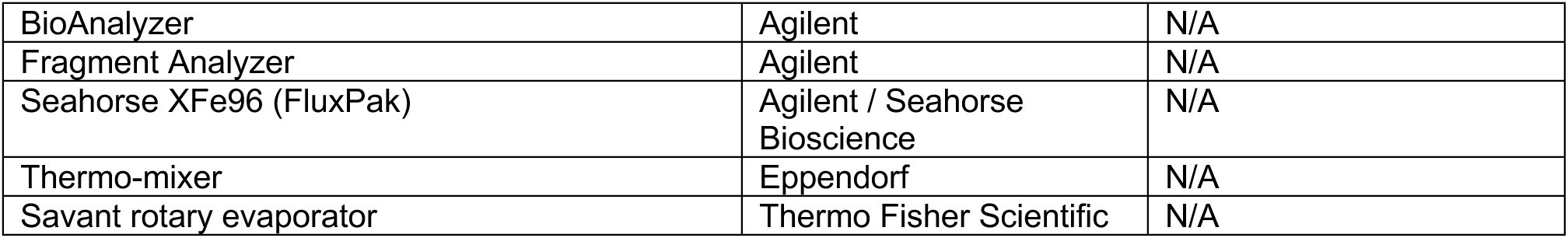

---

**Supplementary Table 1:**
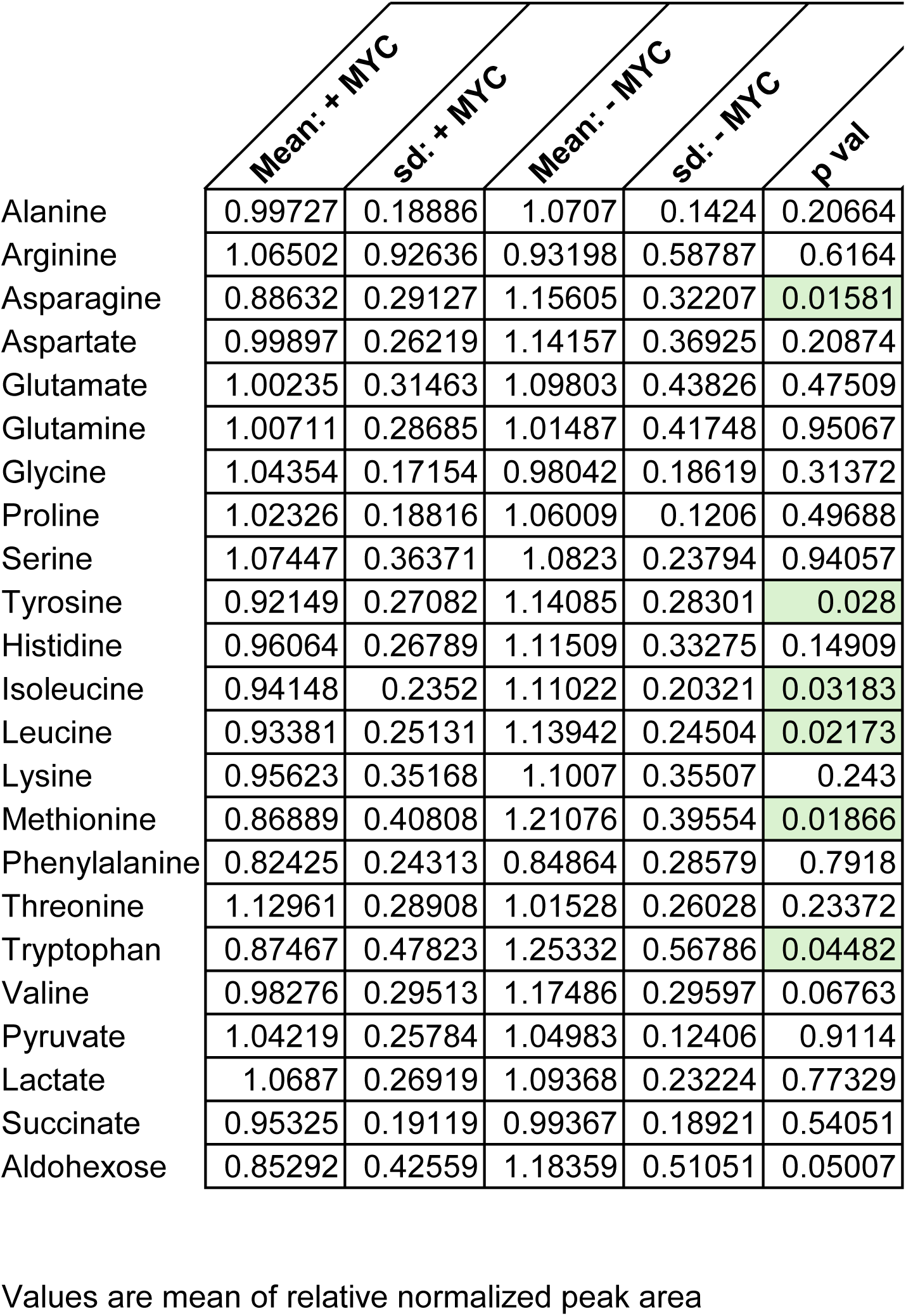
LC-MS analysis of ISF.

**Supplementary Table 2:** LC-MS analysis of plasma.

|  | Mean: +MYC | sd: +MYC | Mean: -MYC | sd: -MYC | Mean: normal | sd: normal | p value (+MYC/-MYC) | p value (+MYC/WT) | p value (-MYC/WT) |
| --- | --- | --- | --- | --- | --- | --- | --- | --- | --- |
| Alanine | 0.852 | 0.251 | 1.022 | 0.188 | 1.157 | 0.318 | 0.042 | 0.007 | 0.171 |
| Arginine | 0.859 | 0.425 | 0.942 | 0.391 | 1.24 | 0.595 | 0.577 | 0.051 | 0.12 |
| Asparagine | 0.821 | 0.297 | 1.021 | 0.26 | 1.189 | 0.345 | 0.055 | 0.004 | 0.15 |
| Aspartate | 1.136 | 0.619 | 0.986 | 0.441 | 0.85 | 0.194 | 0.447 | 0.109 | 0.298 |
| Glutamate | 0.869 | 0.411 | 1.119 | 0.451 | 1.031 | 0.308 | 0.116 | 0.238 | 0.545 |
| Glutamine | 0.888 | 0.196 | 1.016 | 0.128 | 1.123 | 0.205 | 0.041 | 0.003 | 0.102 |
| Glycine | 0.906 | 0.225 | 1.017 | 0.154 | 1.097 | 0.199 | 0.12 | 0.021 | 0.235 |
| Proline | 0.852 | 0.243 | 1.017 | 0.187 | 1.172 | 0.193 | 0.044 | 5E-04 | 0.037 |
| Serine | 0.829 | 0.23 | 1.015 | 0.205 | 1.181 | 0.353 | 0.024 | 0.003 | 0.13 |
| Tyrosine | 0.849 | 0.259 | 1.14 | 0.341 | 1.035 | 0.223 | 0.012 | 0.046 | 0.338 |
| Histidine | 0.883 | 0.231 | 1.062 | 0.165 | 1.07 | 0.221 | 0.02 | 0.033 | 0.911 |
| Isoleucine | 0.857 | 0.254 | 1.101 | 0.164 | 1.052 | 0.232 | 0.004 | 0.038 | 0.511 |
| Leucine | 0.839 | 0.276 | 1.136 | 0.202 | 1.039 | 0.25 | 0.002 | 0.048 | 0.263 |
| Lysine | 0.853 | 0.25 | 1.051 | 0.184 | 1.12 | 0.236 | 0.018 | 0.006 | 0.385 |
| Methionine | 0.805 | 0.269 | 1.109 | 0.257 | 1.111 | 0.242 | 0.003 | 0.003 | 0.981 |
| Phenylalanine | 0.899 | 0.253 | 1.056 | 0.159 | 1.056 | 0.24 | 0.05 | 0.094 | 0.997 |
| Threonine | 0.913 | 0.246 | 0.999 | 0.166 | 1.11 | 0.234 | 0.266 | 0.034 | 0.151 |
| Tryptophan | 0.843 | 0.35 | 1.117 | 0.242 | 1.057 | 0.25 | 0.017 | 0.068 | 0.512 |
| Valine | 0.852 | 0.25 | 1.076 | 0.201 | 1.094 | 0.261 | 0.01 | 0.015 | 0.836 |

Values are mean of relative normalized peak area.

**Supplementary Table 3:** LC-MS of amino acid levels in KPC cells.

| Metabolite | Mean ctrl | Mean DOX | Mean ctrl+CQ | Mean DOX+CQ | sd ctrl | sd DOX | sd ctrl+CQ | sd DOX+CQ | p value(DOX/ctrl) | p value(DOX+CQ/DOX) | p value(DOX+CQ/+ctrl+CQ) |
| --- | --- | --- | --- | --- | --- | --- | --- | --- | --- | --- | --- |
| Asparagine | 1 | 1.3652 | 0.9957 | 1.1014 | 0.074 | 0.0502 | 0.1896 | 0.0333 | 0.0021 | 0.0016 | 0.3953 |
| Methionine | 1 | 2.2499 | 0.5167 | 0.7849 | 0.0655 | 0.5453 | 0.0482 | 0.0384 | 0.0169 | 0.0097 | 0.0017 |
| Alanine | 1 | 1.4656 | 1.398 | 1.7364 | 0.0674 | 0.1074 | 0.3393 | 0.1465 | 0.0031 | 0.0612 | 0.1879 |
| Histidine | 1 | 1.5538 | 0.8293 | 0.9125 | 0.188 | 0.1711 | 0.1617 | 0.0507 | 0.0195 | 0.0034 | 0.443 |
| Isoleucine | 1 | 1.7031 | 0.9624 | 1.2389 | 0.1195 | 0.1873 | 0.1979 | 0.0158 | 0.0054 | 0.0129 | 0.0734 |
| Aspartate | 1 | 1.3973 | 0.8789 | 0.8444 | 0.1031 | 0.1714 | 0.127 | 0.041 | 0.0263 | 0.0056 | 0.6774 |
| Leucine | 1 | 1.9292 | 0.7487 | 0.9699 | 0.0959 | 0.3433 | 0.1392 | 0.0633 | 0.0107 | 0.0089 | 0.0664 |
| Serine | 1 | 1.4436 | 1.0531 | 1.364 | 0.1073 | 0.1279 | 0.1708 | 0.0537 | 0.01 | 0.3766 | 0.0396 |
| Valine | 1 | 1.6319 | 1.0666 | 1.1936 | 0.1696 | 0.1846 | 0.2873 | 0.0818 | 0.012 | 0.0198 | 0.5022 |
| Phenylalanine | 1 | 1.5137 | 1.0574 | 1.102 | 0.1595 | 0.19 | 0.2657 | 0.0524 | 0.023 | 0.0224 | 0.7894 |
| Glutamine | 1 | 2.6383 | 0.7357 | 1.3865 | 0.1167 | 0.9101 | 0.1148 | 0.4961 | 0.0365 | 0.1046 | 0.0913 |
| Glutamate | 1 | 1.2326 | 0.8907 | 1.0463 | 0.0877 | 0.111 | 0.158 | 0.0369 | 0.0464 | 0.0508 | 0.1719 |
| Arginine | 1 | 1.814 | 0.7877 | 1.1027 | 0.2767 | 0.4106 | 0.1457 | 0.2461 | 0.0465 | 0.0618 | 0.1291 |
| Tyrosine | 1 | 1.5274 | 1.5633 | 1.711 | 0.1354 | 0.3237 | 0.3253 | 0.2999 | 0.0598 | 0.511 | 0.594 |
| Tryptophan | 1 | 1.6673 | 1.4471 | 1.5135 | 0.1599 | 0.4528 | 0.4213 | 0.2882 | 0.0738 | 0.6458 | 0.8327 |
| Glycine | 1 | 1.0565 | 1.419 | 1.3492 | 0.1208 | 0.1209 | 0.4783 | 0.1801 | 0.5976 | 0.0797 | 0.8245 |
| Lysine | 1 | 1.8671 | 0.9142 | 1.3402 | 0.3244 | 0.5773 | 0.274 | 0.3814 | 0.0859 | 0.2576 | 0.1912 |
| Proline | 1 | 1.1718 | 1.0763 | 1.0628 | 0.1315 | 0.0749 | 0.235 | 0.0482 | 0.1207 | 0.1013 | 0.9269 |
| Threonine | 1 | 1.2644 | 1.4788 | 1.4395 | 0.1083 | 0.2 | 0.3741 | 0.1184 | 0.1144 | 0.2619 | 0.8708 |

Values are peak area normalized to control cells.

**Supplementary Table 4:**
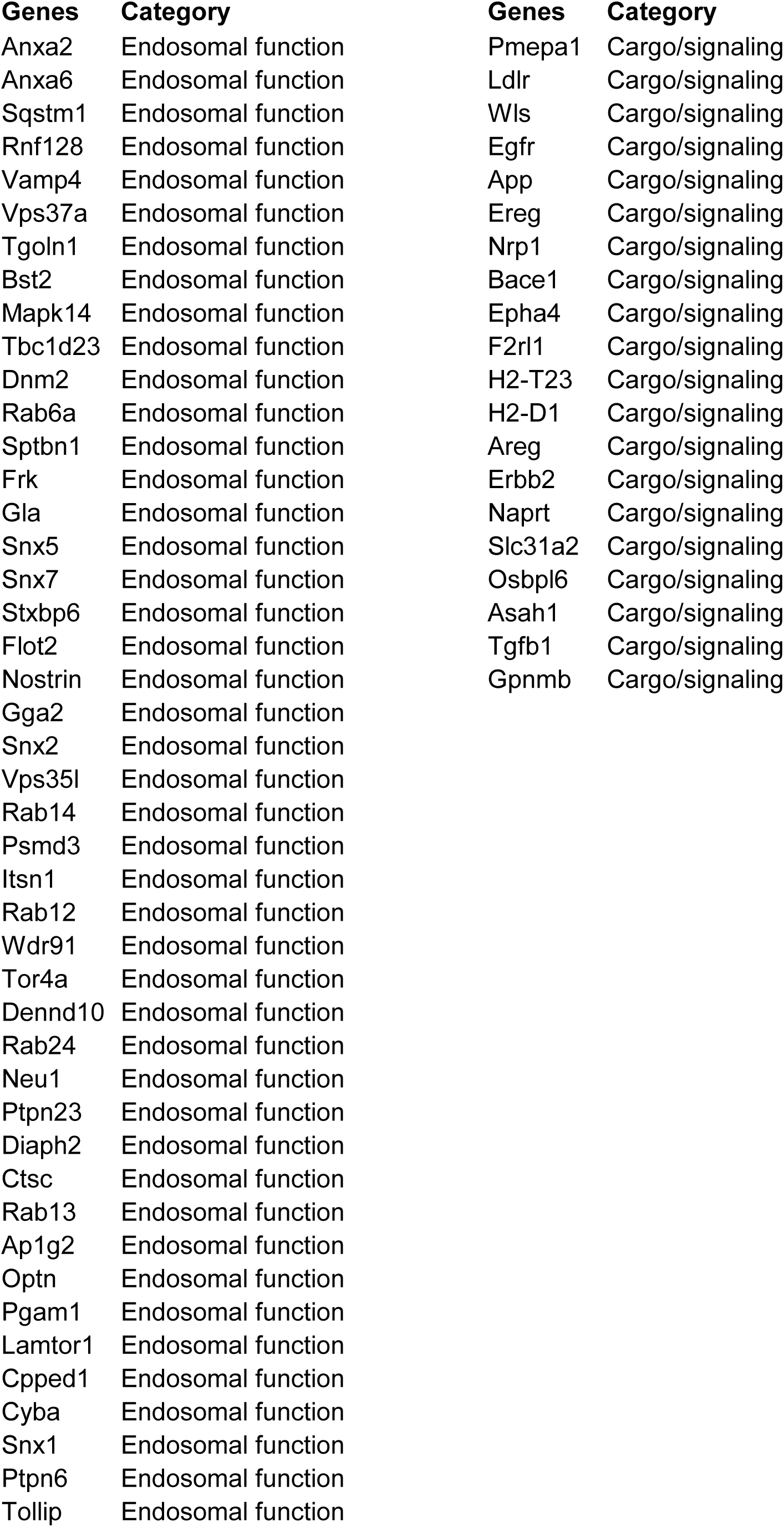
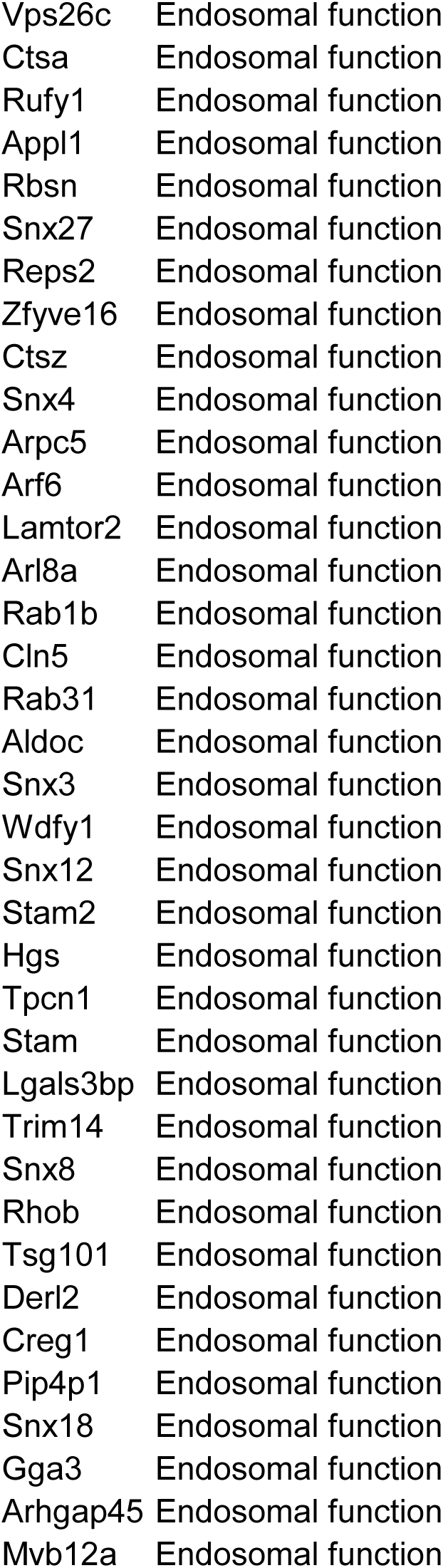
List of gene symbols of proteins from endosomal compartments that are significantly different in SILAC proteomics, categorized into endosomal function and cargo/signaling proteins.

**Supplementary Table 5:** Modeling parameters for the simulation in Supp Data 1.

| symbol | meaning | value |
| --- | --- | --- |
| $s_0$ | baseline survival per generation | 0.8 |
| c | dosage cost per ecDNA copy (linear) | 0.01 |
| a_self | max cell-autonomous protection | 0.9 |
| a_pub | max bystander protection | 0.9 |
| Kp | self half-saturation (own copies) | 4 |
| Ks | bystander half-saturation ($\sigma$) | 20 |
| d_imm | Immune killing rate, unprotected | 0.4 |

**Condition-defining switches**, the one term that changes per panel
| panel | modification |
| --- | --- |
| 1 · CD4-depleted | d_imm = 0 |
| 2 · bystander OFF | a_pub = 0 |
| 3 · full model | (none) |

**Simulation settings**
|  | value |
| --- | --- |
| lattice | 81 × 81, Moore-8, |
| founder | 15 cells |
| founder ecDNA | skewed, mean 2, capped at 8, uniform dist |
| generations | 35 |
| protection form | independent risks (toggle option - additive risk) |
| RNG | mulberry32 (seeded) |
| Seed used in fig 7E | 580 |

Note: Can be changed in each simulation run

